# GeneNetwork: a continuously updated tool for systems genetics analyses

**DOI:** 10.1101/2020.12.23.424047

**Authors:** Pamela M. Watson, David G. Ashbrook

## Abstract

GeneNetwork and its earlier iteration, WebQTL, have now been an important database and toolkit for quantitative trait genetics research for two decades. Recent improvements to GeneNetwork have reinvigorated it, including the addition of data from 10 species, multi-omics analysis, updated code, and new tools. The new GeneNetwork is now an exciting resource for predictive medicine and systems genetics, which is constantly being maintained and improved. Here, we give a brief overview of the process for carrying out some of the most common functions on GeneNetwork, as a gateway to deeper analyses, demonstrating how a small number of plausible candidate genes can be found for a typical immune phenotype.

## Introduction

It is abundantly clear that no aspect of biology works in isolation: cells are filled with networks of interacting proteins; the environment interacts constantly with gene expression; and seemingly tiny perturbations of the genome can lead to lethal diseases. Systems genetics seeks to investigate this by integrating all levels of biology. To do this requires coherent data, gathered together into an easily accessible format, not siloed into disparate data pools that cannot easily be integrated, validated, or extended. This approach will allow us to make animal models of so called ‘precision’ medicine, although perhaps more accurately, we want predictive medicine, where a phenotypic outcome (such as disease) can be predicted, and avoided.

***GeneNetwork*** (genenetwork.org; GN) is one tool for systems genetics and predictive medicine, and this newest version of the web service is the latest iteration of the website that started in 2001 as *WebQTL* (Chesler et al., 2003, 2004). GN provides researchers access to large amounts of diverse data sets ranging from molecular profiles to classical phenotypes, in both a genetically and computationally coherent manner (Sloan et al., 2016; Wang et al., 2016; Mulligan et al., 2017; Parker et al., 2017; Li et al., 2018). Although GN started life as a repository for CNS morphometric and behavioral data for the BXD family of mice, it has expanded to include many datasets, across a variety of species and designs, as well as a toolkit for their analysis. A simple Google Scholar search for ‘GeneNetwork.org’ or ‘WebQTL’ shows over 1400 papers have referenced this resource.

The traditional use of GN was for ‘simple’ forward genetics: identifying a genetic locus underlying variation in a phenotype of interest. However, it has grown far beyond this, with the addition of extensive, coherent datasets, allowing for the integration of multiple levels of biology, as data gathered in different laboratories at different times can be seamlessly combined to provide novel insight. As whole genome-sequence data has become available for many populations used in GeneNetwork, reverse genetics has become possible: moving from variants or genetic loci of interest, and identifying the phenotypes that they influence, so called phenome-wide association studies (PheWAS) (Denny et al., 2010) or reverse complex trait analysis (Li, Mulligan et al., 2010).

GeneNetwork provides to tools and data to do forward, reverse, and systems genetics. Exploring genes, molecules, and phenotypes is easily accomplished using GeneNetwork. In this manuscript we will outline some simple use cases, and show how a small number of plausible candidate genes can be identified for an immune phenotype.

### 1. Data

Once you have navigated to genenetwork.org, there are two ways to search for data in GN. The first is to use the global search bar located at the top of the page (**Figure 1)**. This is a new feature in GN that allows researchers to search for genes, mRNAs, or proteins across all of the datasets. This will give the user data for that search term across many different species, groups, and types of data. Because of this, the global search bar is a good area to start one’s searches if you have a particular gene of interest, but are unsure of what species or populations are relevant, or simply to get an overview of what datasets are available on GN. When searching for genes, it is critical to use standard gene symbols containing more than two characters. This search takes a few minutes to complete, as it is searching through several hundred data sets across species with millions of traits. Although this search will return with a large collection of results, up to 6000 rows, it is also simple to filter on the results page. This demonstrates one of the highlights of GN: large omics datasets have already been analyzed and checked for errors, and then formatted so that they can be directly compared. These seemingly simple points can often take up a large amount of a bioinformatician’s time, and make datasets inaccessible to ‘wet-lab’ scientists.

**Figure 1:**
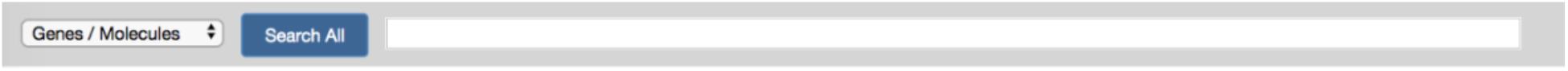
The global search bar, also called the *Search All* function, is a good area to start exploring genes, mRNA, and proteins within GeneNetwork. To best use this new tool, use standard gene symbols containing more than two characters in the name.

Similarly, by using the dropdown menu on the left (**Figure 1**), a user can switch to phenotypes, and search for any phenotype of interest in the same way.

Another area to acquire data is the ***Select and search*** pull-down menus (**Figure 2)**. To get started, the user has to choose a population of interest. GeneNetwork contains data from a wide range of species, from humans to soybeans, but most of the available phenotypic data is from mice. Within the mouse dataset there are groups of families, crosses, non-genetic groupings, and individual data. The type of dataset must be selected after defining the species and sample population. While genotypes, mRNA, methylated DNA, protein, metagenomic, and metabolome datasets are available (i.e. molecular omics phenotypes), the simplest starting point is the phenotype dataset.

**Figure 2:**
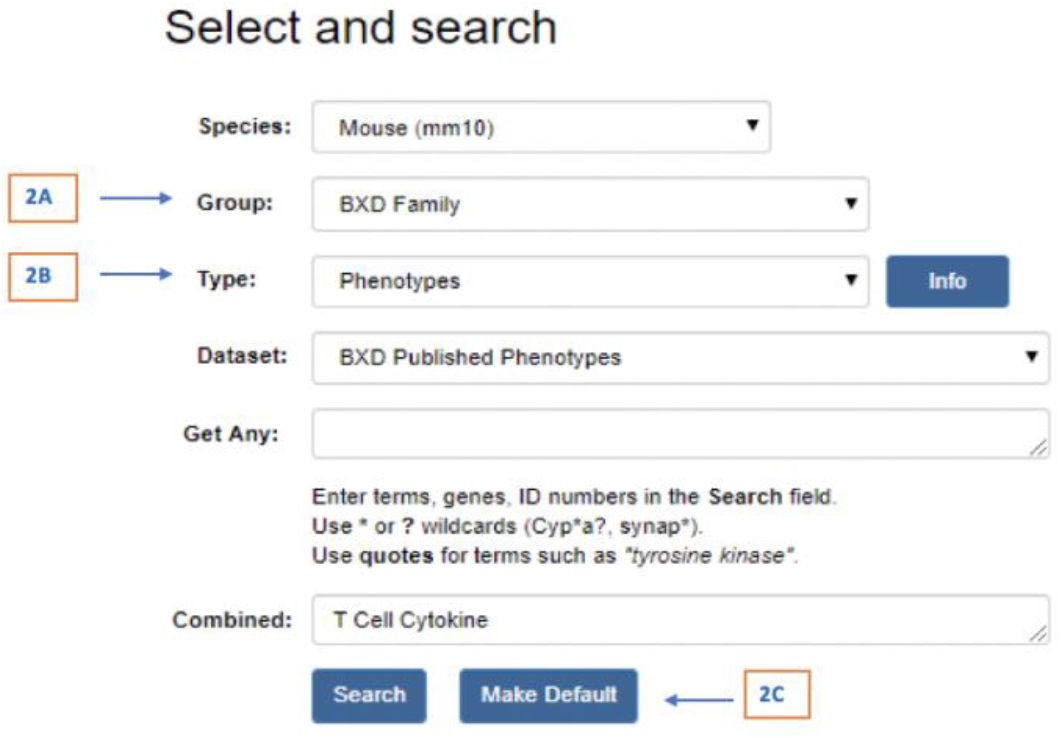
***Select and search*** menus make it simple to access a database of phenotype, omics, and genotype data for humans and plant and animal populations and families. For the purposes of this primer, the ***Mouse BXD family*** (2A) ***Phenotypes*** (2B) is selected. The *Make Default* button (2C) allows users to ‘lock in’ their choice of search criteria, immediately returning to the same dataset the next time GN is opened.

We will use the BXD family (**Figure 2A**)(Ashbrook et al., 2019), for this demonstration, as it has the largest phenome group of over 7000 phenotypes collected over 40 years (**Figure 2B**), as well as over 100 omics datasets, allowing the broadest systems genetics analysis. They have the advantage of each genome being infinitely replicable, unlike F2 or outbred populations, where each individual represents a unique genome.

**Figure 2** shows the available input fields using the ***Search*** function. The first search field is for an ‘any’ search, whereas the second is for an ‘all’ search. For example, let us imagine that we are interested in the regulation of the immune system. Once the user has found the appropriate search parameters for their study, they can also lock that selection by choosing the ***Make Default*** button (**Figure 2C**). To gather a broad collection of phenotypes, we could fill the ***Get Any*** with ‘inflammation immunity infection cytokines’ and this will search for any phenotype description which contains the word inflammation OR the word immunity OR the word infection OR the word cytokines (over 240 entries in June 2020). If, on the other hand, we are only interested in cytokines from T cells, we can use the ***Combined*** search box with ‘T cell cytokine’, which will find any phenotype where the description contains the word ‘T’ AND the word ‘cell’ AND the word ‘cytokine’ (12 entries in June 2020).

Taking this second search as our example, the user must decide which of the phenotypes are relevant to their interest. The initial search produces a records table (**Figure 3)**. This table displays the record ID, a brief description of the project or publication, the authors of the dataset, the year it was published, as well as LRS information. The maximum LRS value will indicate if there is a locus within the BXD family of mice which influences that phenotype. The table also indicates where the maximum LRS value is found within the genome, specifically the location of the marker with the strongest linkage to the phenotype. In **Figure 3A**, we have selected the phenotype with the highest LRS value.

**Figure 3:**
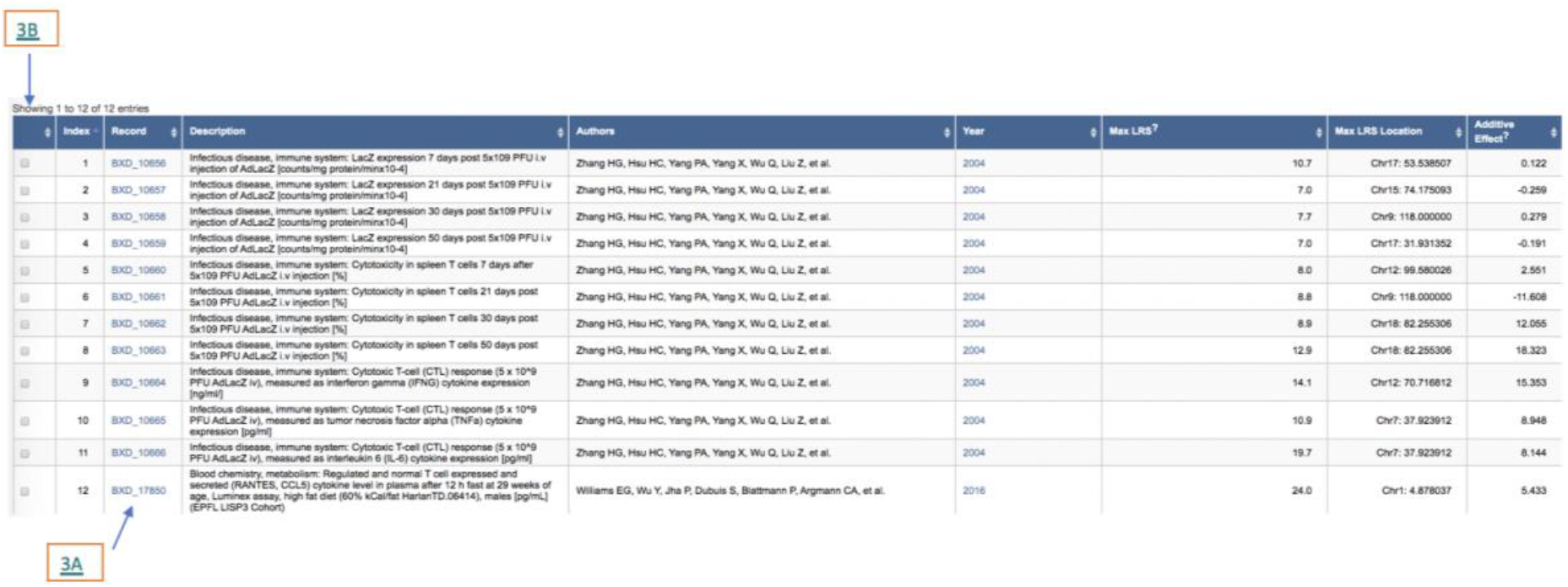
The search results table supplies the necessary information to continue the search for QTLs. Record number, description of the study, authors, year and LRS information are listed to help narrow down possible genetic targets. The table can be sorted by any of these columns, by clicking on the arrow next to the column title. The record ID of interest for this study is 3A, as it has the highest LRS value. 3B highlights the boxes that can be selected in order to form a trait collection.

Another powerful feature of GeneNetwork is the ability to create and analyze whole collections of data. In **Figure 3** there are boxes within the table that can be selected in order to form a trait collection. To do this, select the boxes in the table that suit the interests of the study, and press ***Add***. This function allows groups of traits to be saved for later analysis such as the generation of a QTL, a network graph, and correlation matrix, some of which will be investigated further in this guide.

To access trait data and analysis tools, select the record ID of interest (**Figure 3A**), and this will take you to the record page (**Figure 4**). This record provides a reminder of the species and group being used (**Figure 4A**), and a description of the phenotype (**Figure 4B**). Phenotypes in GeneNetwork ideally follow a set formula, written by the contributor. The first part of the phenotype will be a general grouping, comma separated (Blood chemistry, metabolism). This allows users to easily find large groups of related phenotypes. After the colon is a specific description of the trait, in this case expression levels of the cytokine RANTES (regulated on activation, normal T cell expressed and secreted, also known as CCL5) measured in the plasma using the Luminex assay after 12 hours of fasting, in 29 week old males on high fat diet (Williams et al., 2016). The authors of any related manuscript (or the lab group who gathered the data) are shown, as well as the title and links to the published paper (**Figure 4C**). There is also a button to add the trait to a collection (see below; **Figure 4D**), and to view this trait in the earlier version of GeneNetwork, GN1 (**Figure 4E**). Some less used tools have not been ported across to the latest update, or some users may wish to use a pipeline that they have developed in the past.

**Figure 4:**
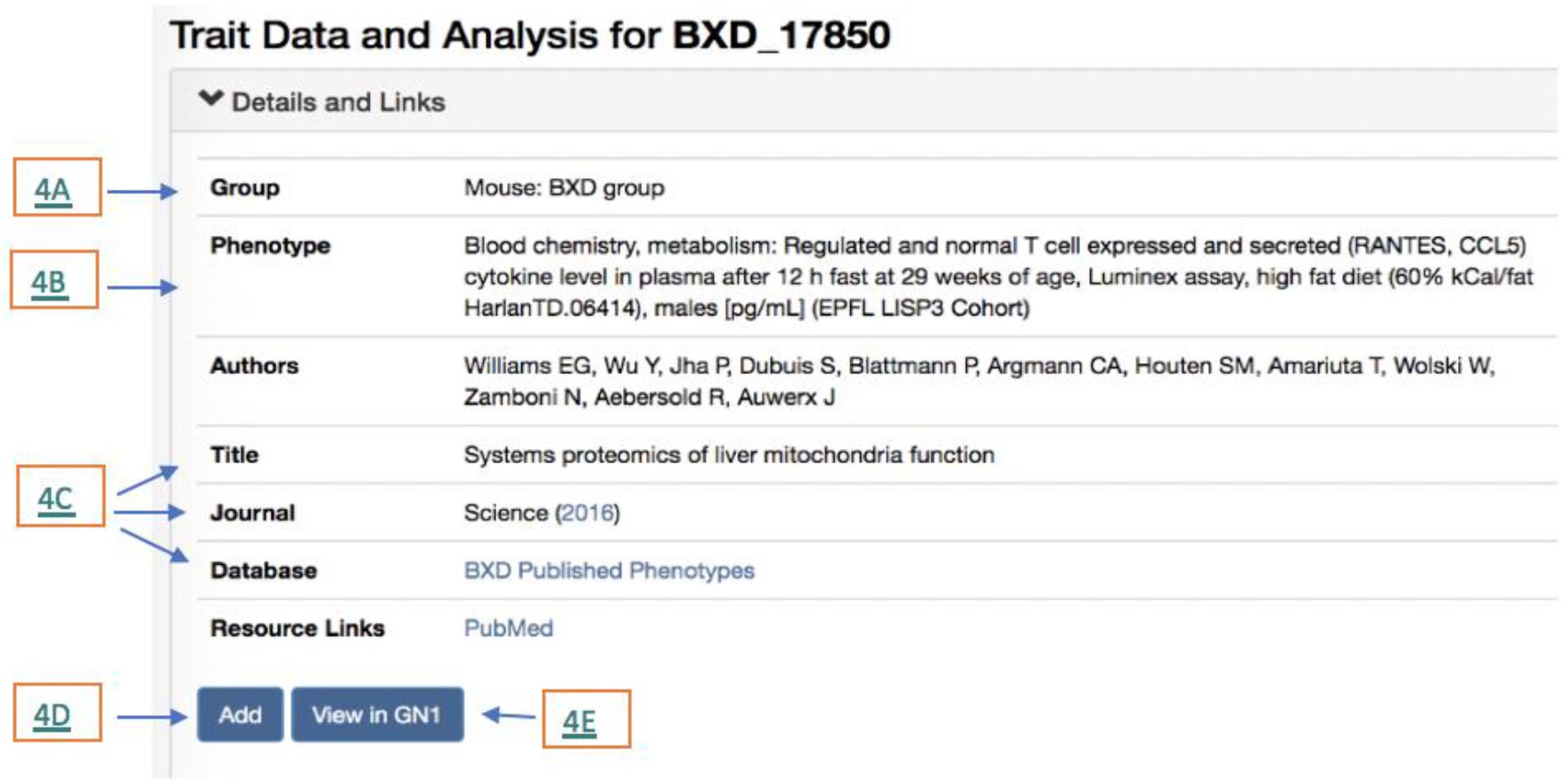
Trait data shows the identifying information of the record including the group name (4A), phenotype being tested (4B), authors of the project, title of the project or paper, journal of publication, and links to the dataset database and to the published paper (4C). There is also an option to add this trait to your collection by pressing the *Add* button (4D), or to view this trait in an earlier version of GeneNetwork, GN1 (4E).

The data uploaded to GN are farther down the page, along with statistical tools to normalize the dataset. The probability plot is a useful tool to visualize the data, as it allows researchers to see the distribution of the data before analyzing it (**Figure 5**). In cases where the distribution is close to normal, the trait values (Y-axis), will correlate well with the theoretical quantiles (X-axis). Samples far away from this X=Y lines are likely outliers, which can skew the results from the analysis. Other deviations, such as an S-shape (somewhat seen in **Figure 5**), abrupt breaks between samples, or a set of ripples, are good indicators that there is one or more large effect QTL for that trait.

**Figure 5:**
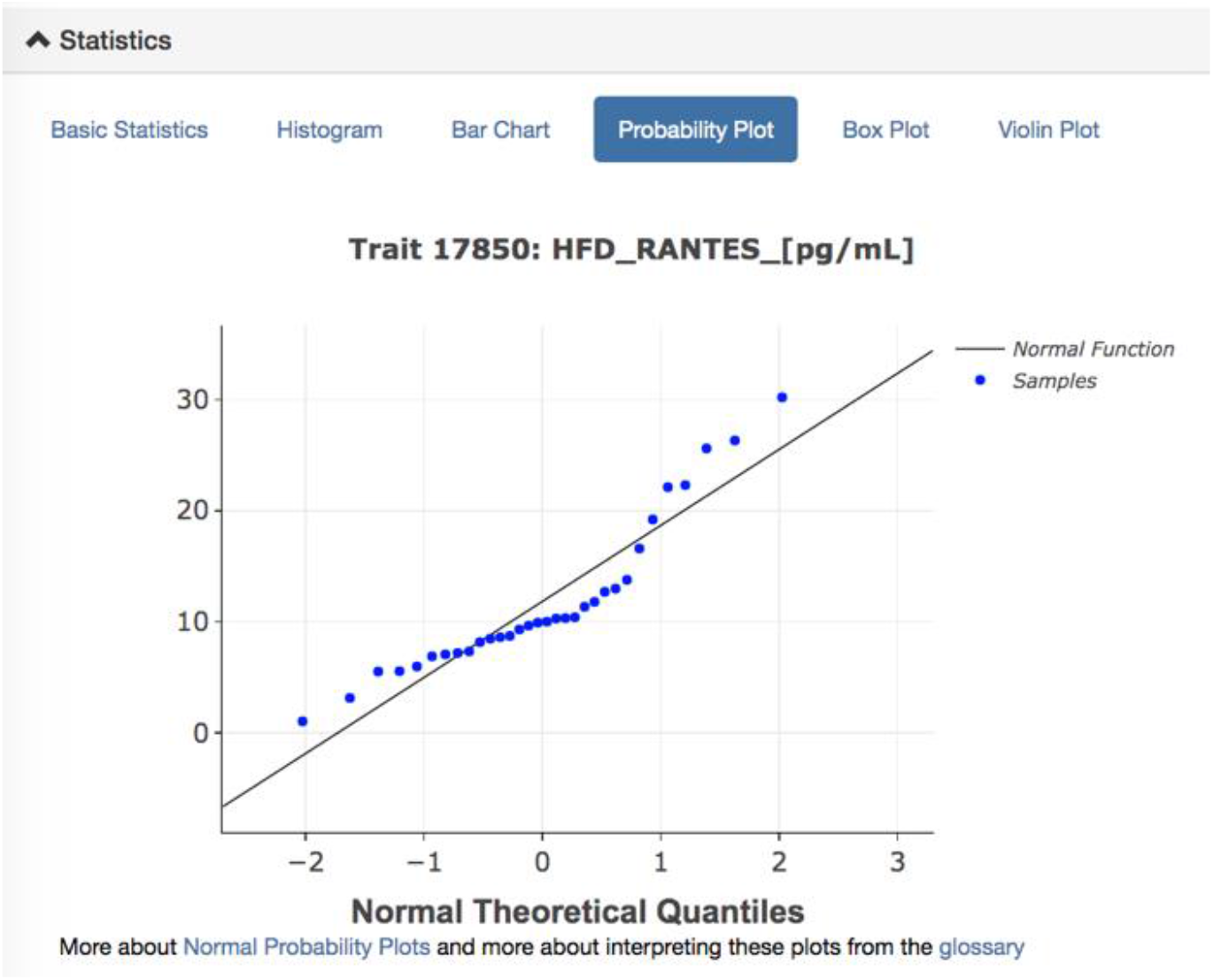
Statistical analysis tools to help normalize the data before investigating QTLs. One of the best ways to initially view the data is by using the *Probability Plot.* This provides viewers with a visual representation of the data. If the data is skewed, it will not follow the normal function line. For this dataset, the probability plot shows that the data is not normal, but is in an S shape. This could suggest that there is a large QTL that could possibly account for that, or that the data needs to be normalized. After viewing, it is then possible to normalize the data with the *Statistical Tools* drop down menu.

In this case, the data distribution is fairly normal. However, the transformation and filtering option is located under the statistical tools drop down menu if necessary. In this function, outliers can be blocked, and the data can be normalized by Log2, Z Score, quantile, square root, or the data can be inverted. Specific data points can also be blocked using this feature.

The bar chart visualization is another useful tool, as when sorted by value it allows us to see both the distribution of strains, and the standard error of values. We can see that BXD75 has a high value, but also a very large SE. In the new GN we can now go down the trait page and remove BXD75 from the analysis to see if it is having an undue effect on the analysis. In this case the QTL is not significantly altered by removing BXD75, and so the stain was kept for subsequent analyses. To do this, we simply press the green reset button.

### 2. Mapping

#### QTL

There are thousands of ‘classical’ phenotypes publicly available on GeneNetwork (GN). These classical phenotypes are mainly whole-body phenotypes similar to the kind that have been collected and studied for over a century. This is in contrast to molecular phenotypes, such as gene expression or epigenetics. GN allows for the integration of these phenotypes, and even entire phenomes, into a systems genetics analysis. One way to analyze large sets of genetic data is to find a quantitative trait locus, or QTL. A QTL (http://gn1.genenetwork.org/glossary.html#Q) is an area on a chromosome that can contain one or many genes, that is linked to a change in phenotype. After a QTL that is responsible for the apparent variation in phenotype has been identified, one can start studying the genes within that locus to identify the likely causal gene.

Once the data is normalized appropriately (in our case, no normalization was required), the QTL can be mapped. To do this, select the mapping tools drop down window (**Figure 6)**. There are three methods to choose from, GEMMA, Haley-Knott Regression, and R/qtl (**Figure 6)**. Genome-wide Efficient Mixed Model Analysis (GEMMA; github.com/genetics-statistics/GEMMA; (Zhou and Stephens, 2012) is a multivariate linear mixed model mapping tool that is used to map phenotypes with SNPs with a correction for kinship or any other covariate of interest. This ability to account for covariates is highly useful, but also this increases the time taken for computations. However, given the complex relatedness amongst the BXD strains, GEMMA is the best tool to use with the BXD cohorts. GEMMA produces a Manhattan plot with chromosomes on the x-axis and a −log(P) scale on the y-axis. This −log(P) is approximately equal to a LOD score in this population. The Haley-Knott Regression is the classic method, which has been used in GeneNetwork for almost 25 years (Haley and Knott, 1992). Haley-Knott Regression uses multiple regression formulas to analyze the likelihood ratios between traits and phenotypes, and these calculations are much faster than those used in GEMMA. However, it is not recommended for closely related populations or admixed populations, as it does not consider relatedness or kinship between samples, and so is not fully appropriate for many populations. It is still very useful for F2 intercrossed populations and backcrosses. Due to the great speed of calculation, Haley-Knott Regression also allows us to carry out permutations to determine an empirical statistical significance threshold: the default on GN is 2000 permutations, but even 20,000 permutations only take a minute or so. For this reason, Haley-Knott Regression is still our preferred option when mapping large omics datasets (where there can be tens of thousands of traits), and for producing permutation thresholds. The Haley-Knott Regression mapping tool produces a map of the chromosomes on the x-axis and the likelihood ratio statistic on the y-axis, although this can be switched to a LOD score as needed. We also provide the option of mapping with the widely used R/qtl tool (Broman et al., 2003). R/qtl allows five different mapping methods, and therefore can be adapted for specific situations. We plan to have the second version of R/qtl (R/qtl2) integrated into GN soon (Broman et al., 2019). We currently recommend using both Haley-Knott Regression to determine a significance threshold and GEMMA to identify the strength and location of the QTL.

**Figure 6:**
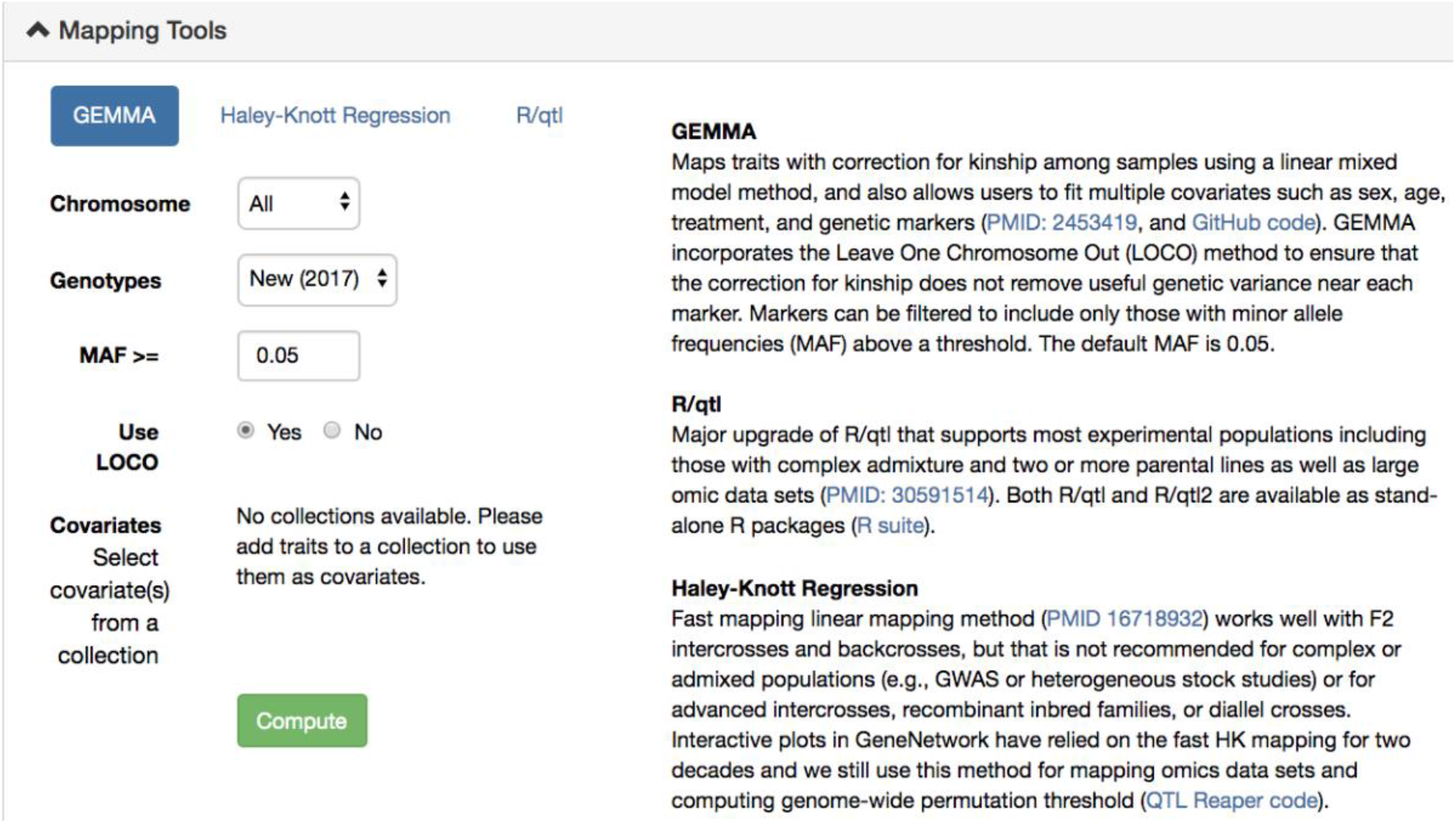
Mapping tools available in GeneNetwork. At the top of the pull-down menu, there are three buttons that can be selected to analyze data using the three mapping tools. We currently recommend using both Haley-Knott Regression and GEMMA together to gather the best results.

After choosing the appropriate mapping method, click compute. The next screen will show LRS scores, suggestive levels, significant levels, and frequency of LRS peak values. The LRS value, the likelihood ratio statistic (http://gn1.genenetwork.org/glossary.html#L), is a value that measures the association between the gene variation and the trait of interest (Parker et al., 2017). The suggestive and significant LRS threshold values are generated by permutations, as mentioned above. The suggestive threshold provides an empirical p-value of 0.63: this level will yield an overage of one false positive per genome-scan. These peaks are worth investigating, for example, if several linked phenotypes have a suggestive locus at the same position, then the position may be significant when the traits are analyzed jointly. The significant threshold is a LRS value with a genome wide empirical p-value of 0.05 (Mulligan et al., 2017). For our trait, the chromosome containing the highest peak is chromosome 1 (**Figure 7)**. In this case, the peak passes the significant LRS value, which means that there is a less than 5% chance that there is not linkage between the phenotype and this locus. Therefore, this locus warrants further investigation.

**Figure 7:**
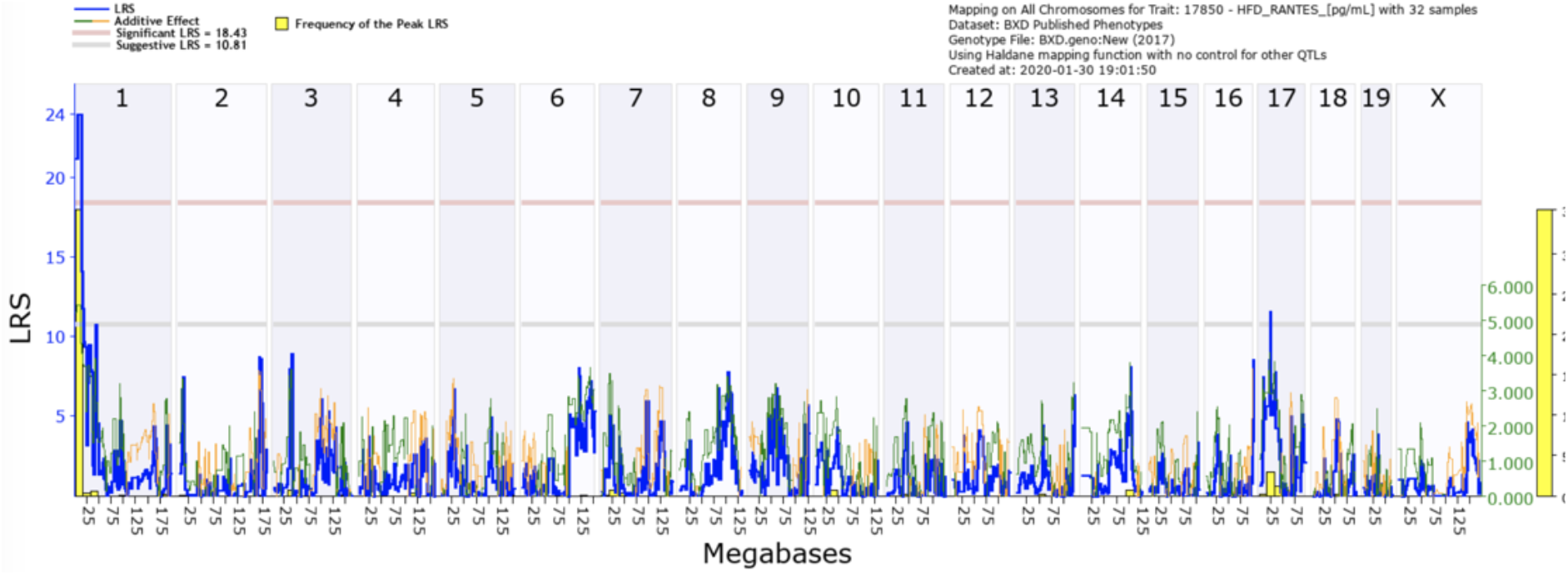
QTL map of the whole genome of the BXD mice in this data set with peaks according to association with the phenotype being tested. The grey line indicates the suggestive LRS values, with a p-value = 0.63 (one false positive per genome scan), as determined by permutations of the data. The red line indicates the significant LRS value (p-value = 0.05), which for this dataset is an LRS value of 18.43. The blue line shows the linkage between the trait and that position along the genome. The orange/green line represents the additive effect. The yellow bar shows the frequency of the LRS peak location from 2000 bootstraps of the data.

We can zoom in to chromosome 1, by clicking on the chromosome number at the top of the QTL map. In this view (**Figure 8)**, individual genes can be observed. The multicolored bars at the top of the figure are individual genes, and the orange dashes on the bottom are the SNP density in that region. The blue line shows the LRS score at that position along the chromosome, and the yellow bar indicates the area with the greatest frequency of highest LRS score in a bootstrap analysis. For a closer look, the red bar on the top of the figure can be selected to zoom in to that region. Similarly, a specific region can be requested by selecting the chromosome and start and end positions at the top of the page. With this focused view, we can now determine the 1.5 LOD drop interval. The 1.5 LOD drop interval is a commonly used threshold to determine the region of the genome with > 95% confidence as containing the causal variant (Manichaikul et al., 2006). LOD = LRS/4.61, and therefore the 1.5 LOD drop threshold is equal to an LRS drop of 6.915. Our peak LRS is 24.026, and so the extent of the locus is determined by the first marker to fall below an LRS value of 17.111. Using this method, we can determine that the 1.5 LOD drop interval runs from the beginning of chromosome 1 to 14.8 Mb.

**Figure 8:**
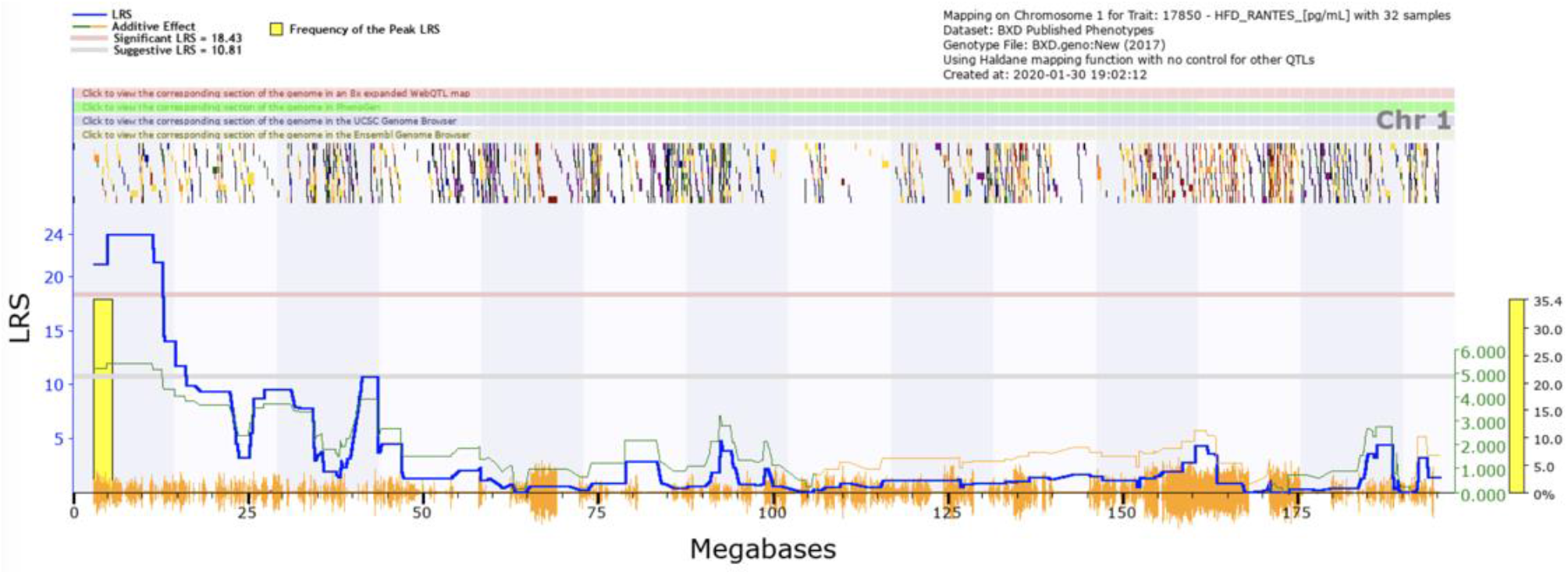
QTL map of chromosome 1, showing a significant QTL between 0 and 25 Mbp, CCL5 expression trait (GeneNetwork ID BXD_17850). The red horizontal line towards the top of the figures illustrates the genome-wide significance threshold, which is p-value ≤ 0.05 genome-wide corrected (significant LRS = 18.43). The lower, blue line indicates the significance of the trait at each position along the genomic regions. The lower, green and red line shows the negative additive coefficient. Regions where DBA/2J alleles increase trait values are shown in green, and regions where the C57BL/6J alleles increase trait values are shown in red, with the scale in green on the left. The multi-colored blocks at the top of the figure represent genes, showing where they fall in the genomic region, and their length. The density of segregating SNPs in the BXD family are shown by the orange seismograph track at the bottom of each map. Adapted from genenetwork.org.

From the zoomed in view of the 1.5 LOD drop interval (**Figure 9**), we can investigate genes, either by selecting the colored bars on the top of the figure or by scrolling down to the interval analyst. This table, shown in **Figure 10**, displays the genetic information for the 96 genes within the mapping interval. Some helpful information within this table is the abbreviation for the gene, the location and length of the gene, the number and density of SNPs in the dataset, the location on the human chromosome, and a description of the gene. Based on the descriptions of genes within the area of interest, the gene *Atp6v1h* was chosen as the most probable candidate for this QTL. This was mainly because of the functions of the gene, specifically the transport of innate immune system components and its involvement in signaling pathways, (Zhang et al., 2017). Selecting the gene opens a new link to the NCBI gene information page, shown in **Figure 11**. This webpage offers summary information for the gene and its functions. This information can assist researchers in discovering genes associated with their phenotypes and QTLs.

**Figure 9:**
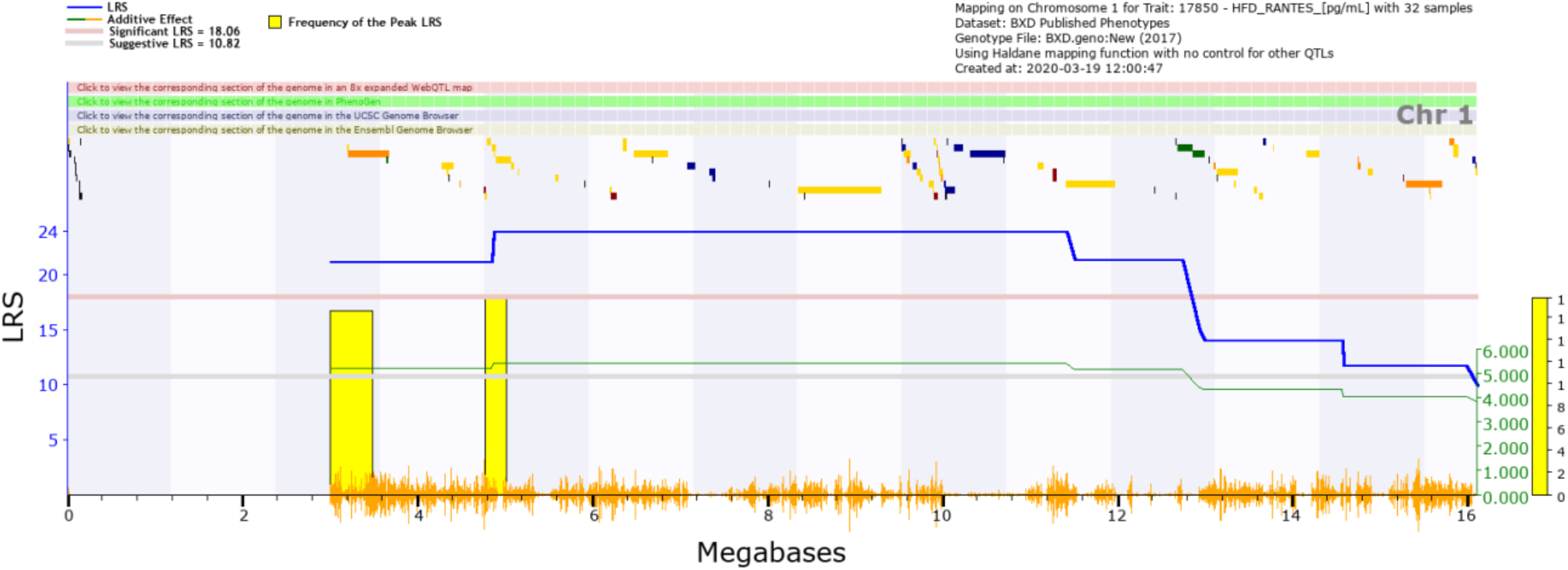
Close up of the 1.5 LOD confidence interval on chromosome 1. The blue line is the LRS value for individual regions on the chromosome. The yellow bar is the frequency of the highest LRS values within the region. The orange hashes on the bottom by the megabase values are to denote areas of high SNP density. The peak LRS value for this data set is 24.06. To calculate the LOD value, take the peak LRS value and divide by 4.61. For this QTL search, the LOD drop interval is anywhere around the peak that is between 24.06 (LOD = 5.21) and 17.111 (LOD = 3.71). This area is found between the beginning of chromosome 1 to about 14.8 Mb.

**Figure 10:**
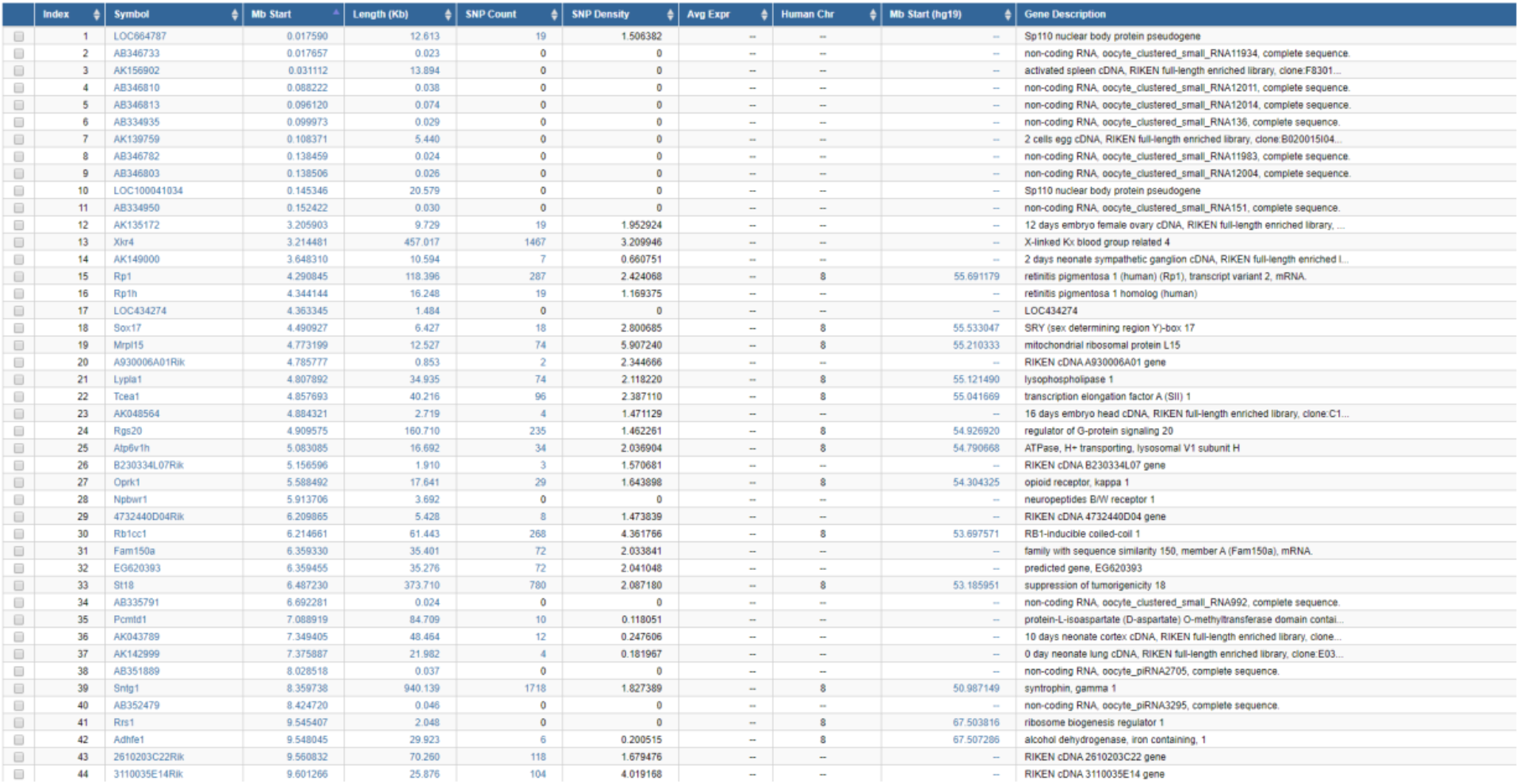
The first 44 lines of a table containing genes within the 1.5 LOD drop confidence interval for our RANTES expression trait (GeneNetwork ID BXD_17850). Information on all genes is given: the current official mouse gene symbol (Symbol); the megabase start position of the gene on the mm9 genome build (Mb Start); the length of the gene in kilobases (Length Kb); the number of SNPs between DBA/2J and C57BL/6J within the gene (SNP Count); the density of SNPs within the gene; the chromosome on which the human homologue is found, if one exists (Human Chr); the megabase position of the human homologue on the hg19 genome build; and the NCBI gene description (Gene Description).

**Figure 11:**
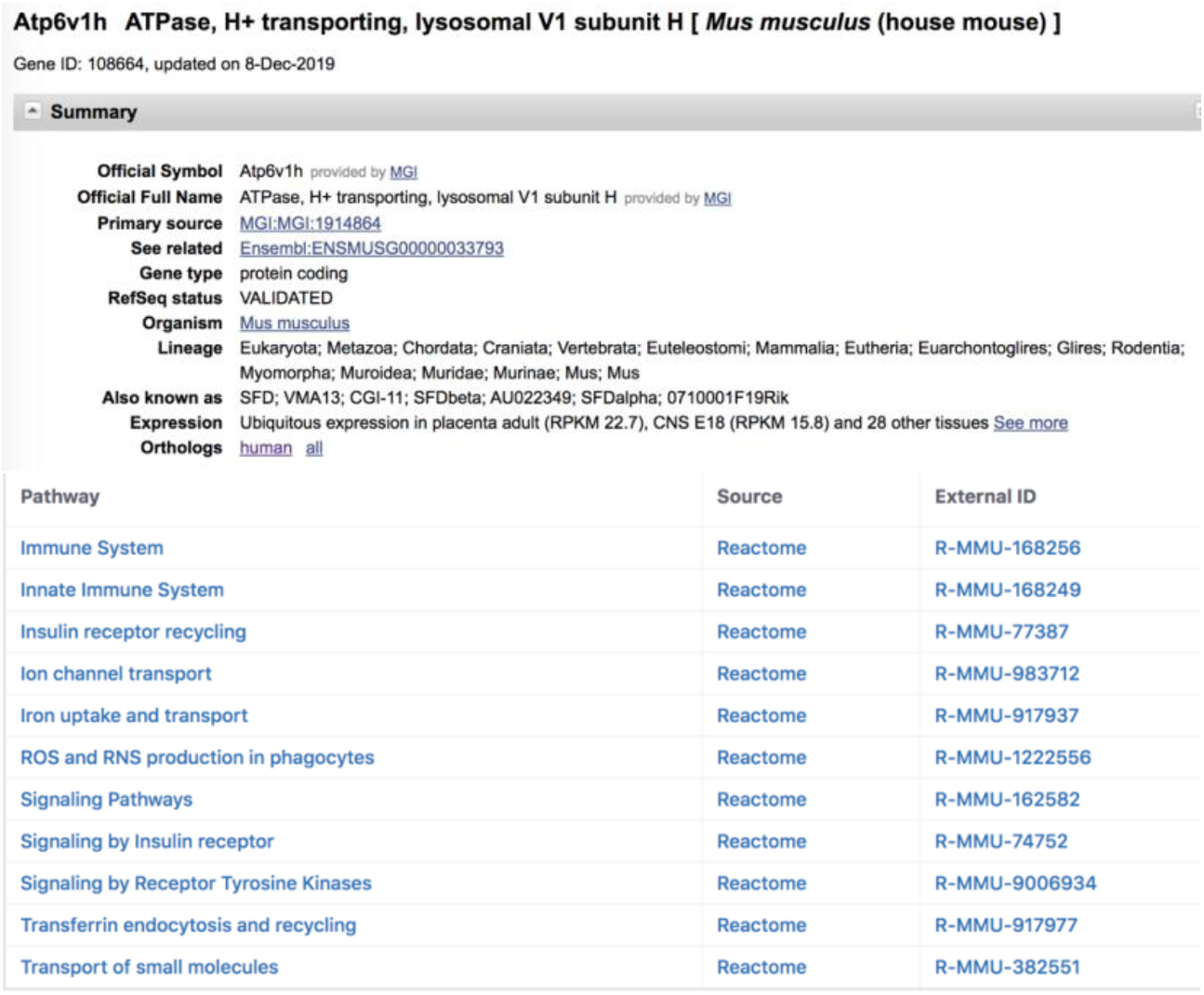
The gene residing within the QTL and its functions as listed within the NCBI database. The summary provides information pertaining to the gene, including its relative expression within the organism of interest. The pathways mentioned on this webpage can offer insight into the gene’s functions. For this investigation, the QTL mapped closely to the *Atp6v1h* gene.

#### Expression QTL

Next, we will examine expression quantitative trait loci (eQTLs). These are QTLs for gene expression traits, a subset of the molecular phenotypes mentioned above. Much like classical phenotypes, expression of transcripts can be influenced by variants within the genome. However, because we know the location of the gene, we can split these eQTL into two categories, *trans-* (or distal) or *cis-* (or local) eQTL.

A *trans*-eQTL (or distal-eQTL) describes when the expression of a gene is influenced by a locus far away from that gene, and therefore indicates that the gene of interest is downstream of another gene. For example, a variant in a transcription factor will alter the expression of its downstream target, which would show a *trans*-eQTL at the position of the transcription factor. This is useful for researchers to know when creating pathways for biological processes or examining genetic interactions.

*Cis*-eQTL describes when a variant within or close to a gene influences its expression, and this can occur when a variant is found in a transcription factor binding site. *Cis*-eQTLs are often of interest as they show that a gene is under its own regulation, and provide us with the beginning of a chain of causality (a variant in that gene causes a change in that genes expression). The overlap between a phenotype QTL and a *cis*-eQTL is strong evidence that the gene is causative of the phenotype.

##### Cis-eQTL

Let us examine *Atp6v1h*, our candidate gene from the first section. From the phenotype we selected, we hypothesize that expression of *Atp6v1h* in T cells is altering the expression of RANTES (regulated on activation, normal T cell expressed and secreted, also known as CCL5). We can next ask the question, does *Atp6v1h* have a *cis*-eQTL in T cells? To accomplish this, go to the home page of GeneNetwork and adjust the search parameters. To match the phenotype, we select the same the ***Species*** and ***Group*** (**Figure 12**). For eQTLs, the ***Type*** of selected data will be one of the ‘mRNA’ datasets, and specifically ‘T cell (helper) mRNA’ (**Figure 12A**). There is only one dataset within this category (HZI Thelp M430v2 (Feb11) RMA) so we will use this (**Figure 12B**). We can then search for Atp6v1h in the ***Get Any*** box (**Figure 12C**).

**Figure 12:**
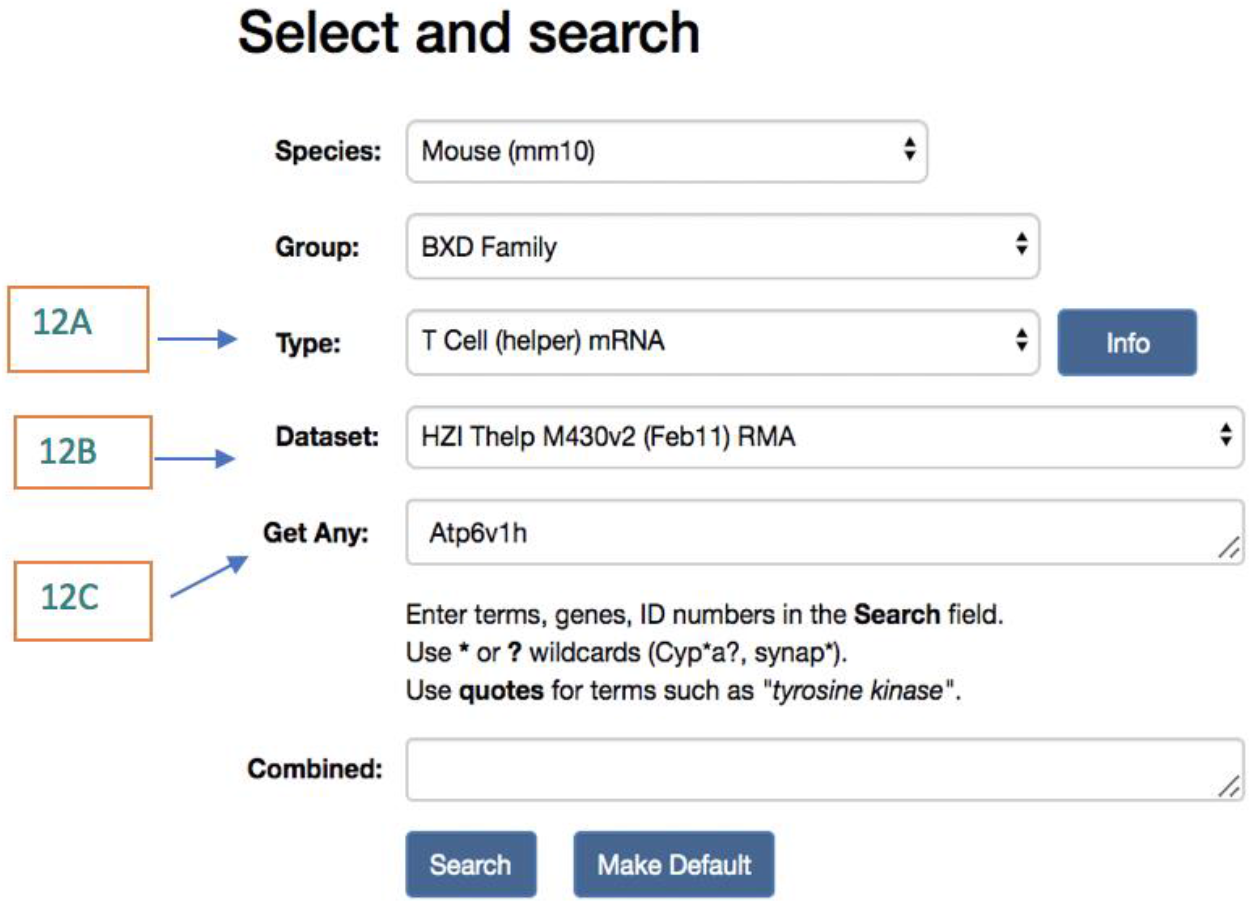
The search parameters on the home page of GeneNetwork for an expression QTL. For this search, the same species and group were used. However, expression is measured by mRNA, so the *Type* (12A) and *Dataset* (12B) are for mRNA. In the *Get Any* box, we input our gene of interest from above (12C).

Just as before, a table with the records matching the search parameters will populate (**Figure 13)**. We can see that there are 5 results, 4 of which are for probes which target *Atp6v1h*. Two of these (1440549_at and 1415826_at) show significant *cis*-eQTL, as the location of the maximum LRS value (***Max LRS)*** is within a few megabases (Mb) on the same chromosome. This is strong evidence that a variant in *Atp6v1h* alters its expression in T-helper cells. If the peak LRS occurs within 10 Mb of the gene of interest, it is considered a *cis*-eQTL, although this distance will depend on the number of recombinations and accuracy of genotyping in a particular population. To further investigate this, select the record named 1440549_at. The *Trait Data and Analysis* table will appear on the next screen, **Figure 14**.

**Figure 13:**
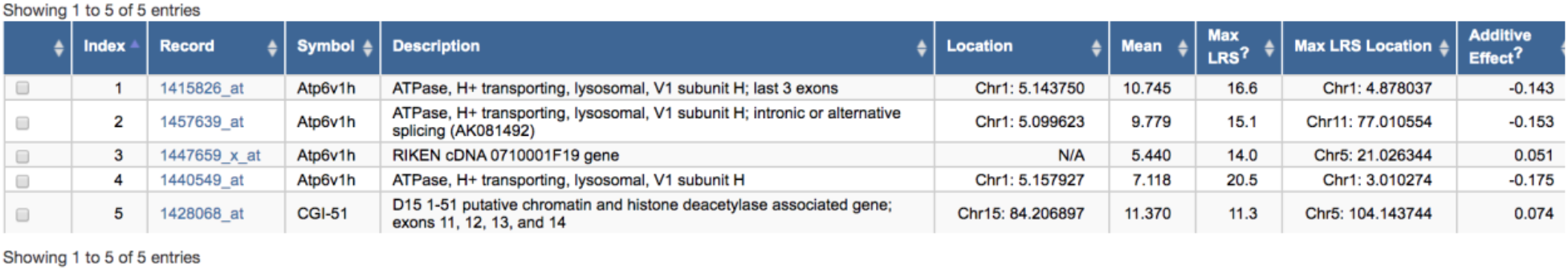
The records table for the eQTL search. 4 of the results are for probes that target the gene of interest, *Atp6v1h*. Of those 4, two show a significant cis-eQTL (1440549_at and 1415826_at).

**Figure 14:**
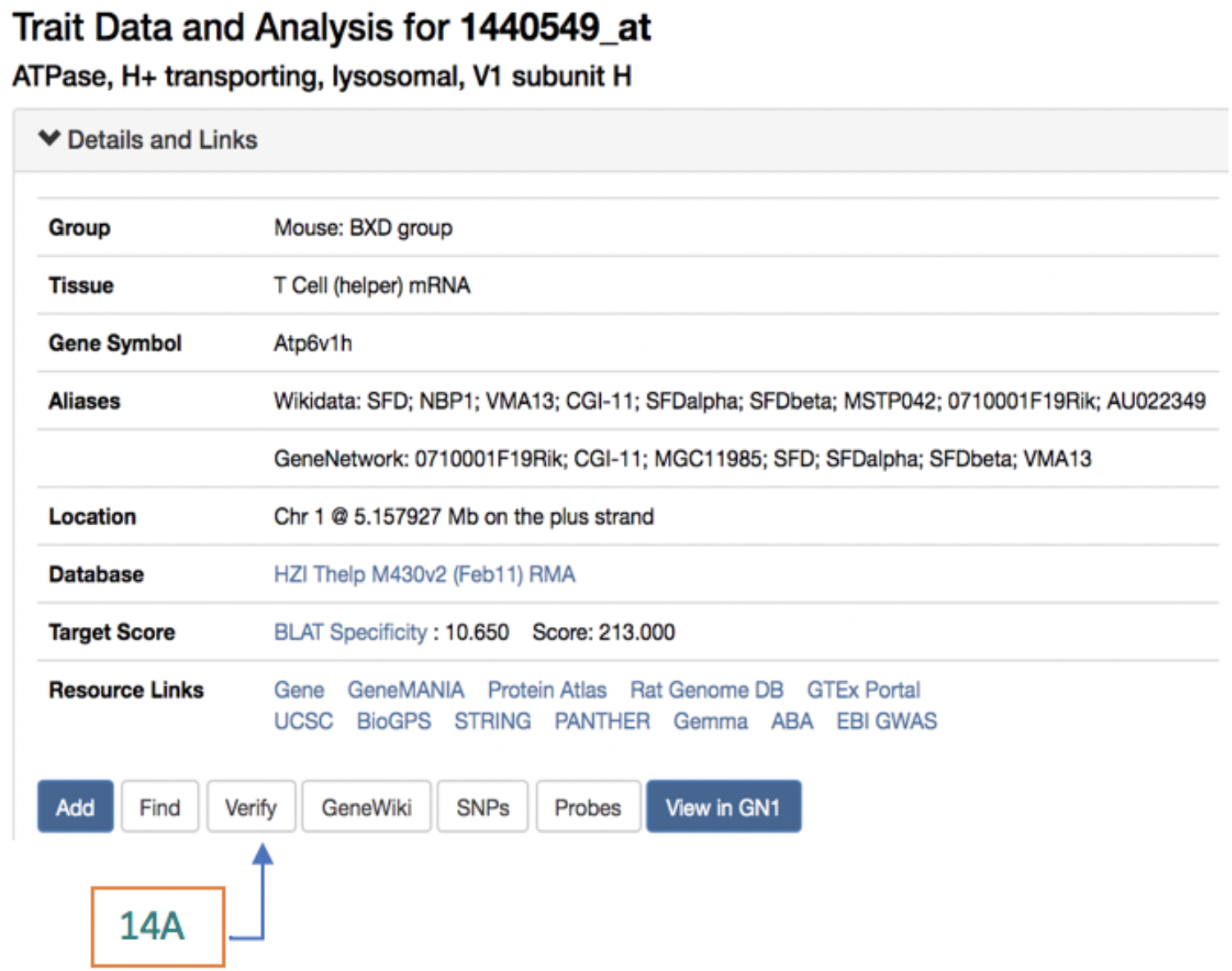
Trait and data analysis for the 1440549_at probe for *Atp6v1h*. The window for the Trait Data and Analysis page looks very similar to the one for the QTL, however there are more buttons below the resource links. An important one to utilize before checking for an eQTL would be the Verify button (14A). This redirects to the UCSC BLAT website to verify the integrity of the probes used for the microarray.

It is important to validate the probe used for the assay. Some probes are inefficient, could accidentally target other mRNAs, or are disrupted by variants in the population. One way to validate the probes used the UCSC BLAT website (Kent, 2002). BLAT is a tool that aligns the probes used to the reference genome to identify off-target alignments. To do this, select the ***Verify*** button on the trait analysis table (**Figure 14A**). This will open a new window that has the probes being verified as well as the alignment scores (**Figure 15)**.

**Figure 15:**
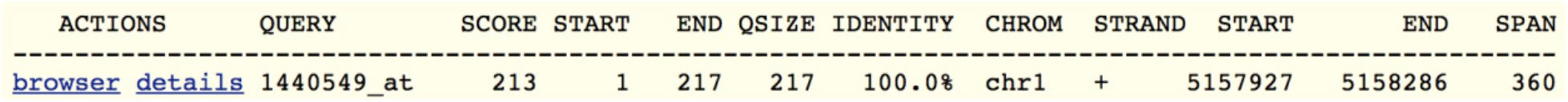
The UCSB BLAT website is one way to verify the probes used to collect the mRNA data. In this case, the probe has 100% identity to the gene of interest when aligned with the reference genome and does not have off target effects for the eQTL search.

Finally, after validating the probe, check the distribution of the data. As before, if the data is not distributed normally, it is essential to normalize before trying to map the location of any QTL. The data for this example was distributed normally, normalization not necessary (**Figure 16**).

**Figure 16:**
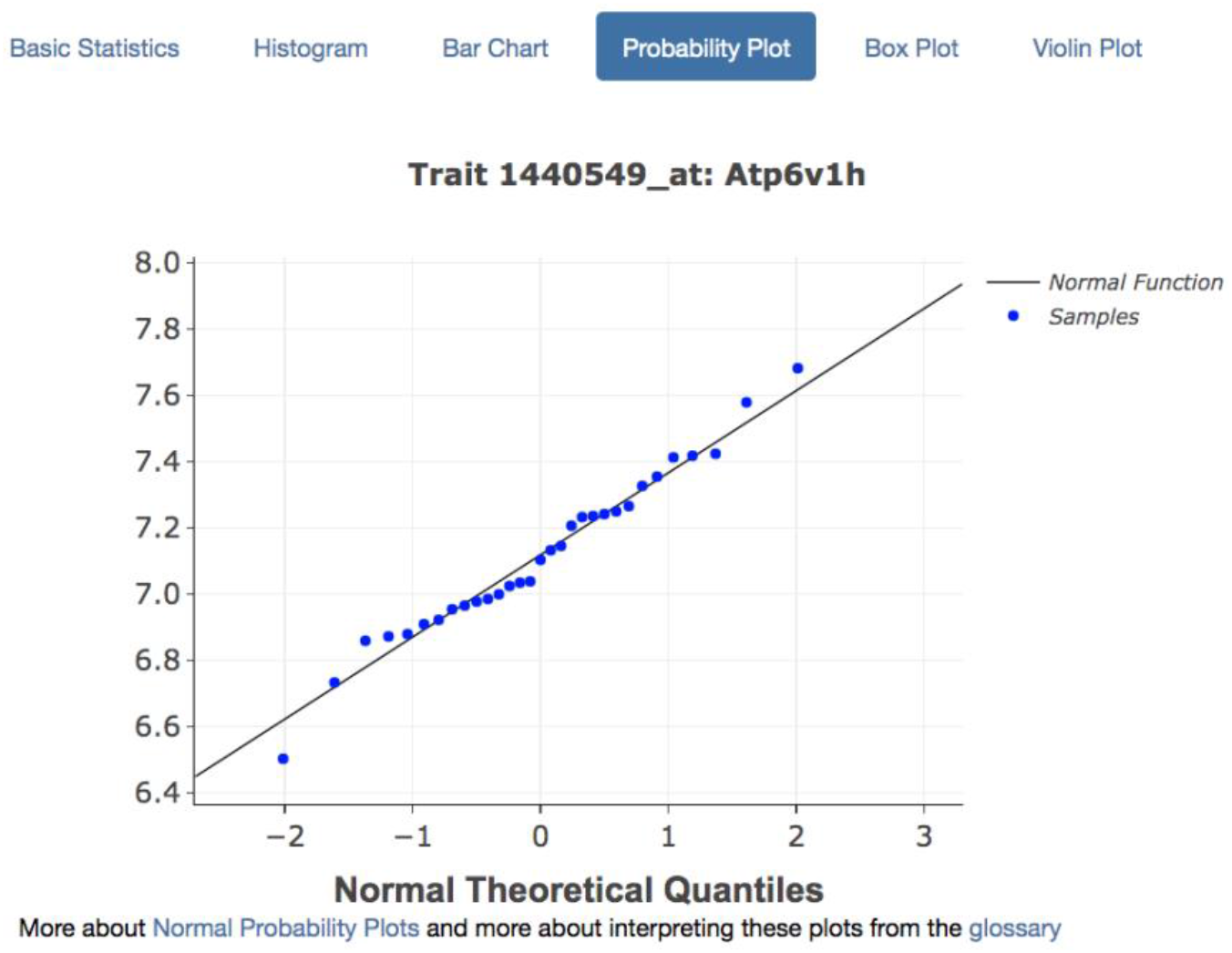
Normalization of eQTL data. The data appears to be normally distributed, so we can proceed with the mapping. However, just as before the tools for normalizing the data are found in the Transform and Filter Data drop-down menu.

After checking the distribution of the data, choose the mapping method relevant to the dataset and select compute. As shown previously, the resulting map will populate with peaks according to the LRS score along the genome. In this case, the highest peak is on chromosome 1, shown in **Figure 17**. Note the purple arrow at the bottom of the figure, which shows the position of the probe in the genome.

**Figure 17:**
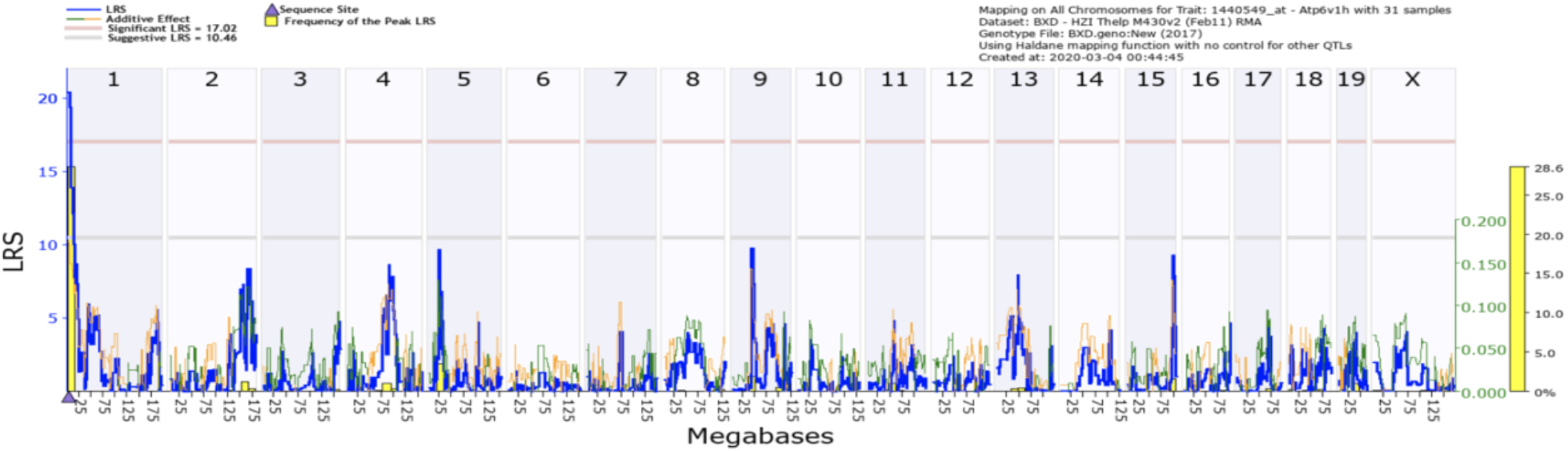
QTL map for the 1440549_at probe for *Atp6v1h*, showing association between expression of the probe and markers across the whole genome of the BXD mice. The grey line indicates the suggestive LRS values, which are those values that have at least an LRS value of 10.81. The red line indicates the significant LRS values (p-value 0.05), which are those values that meet an LRS value of 18.43 for this data set. The blue line shows the LRS value across the genome. The orange/green line represents the additive effect of the B6 (green) or D2 (yellow) allele. The yellow bar shows the frequency of the LRS peak location within the genome from bootstrap analysis.

By clicking on the chromosome, the map is repopulated to include individual genes and SNPs (**Figure 18)**. Again, on the bottom of the map is a purple triangle that marks the placement of the probe for *Atp6v1h*. Upon further expansion of the map, the gene of interest becomes the most likely candidate for its own expression, in that expression of the gene maps back to location of that gene on the genome (**Figure 19**). In this case, the expression of *Atp6v1h* is controlled by a *cis*-eQTL.

**Figure 18:**
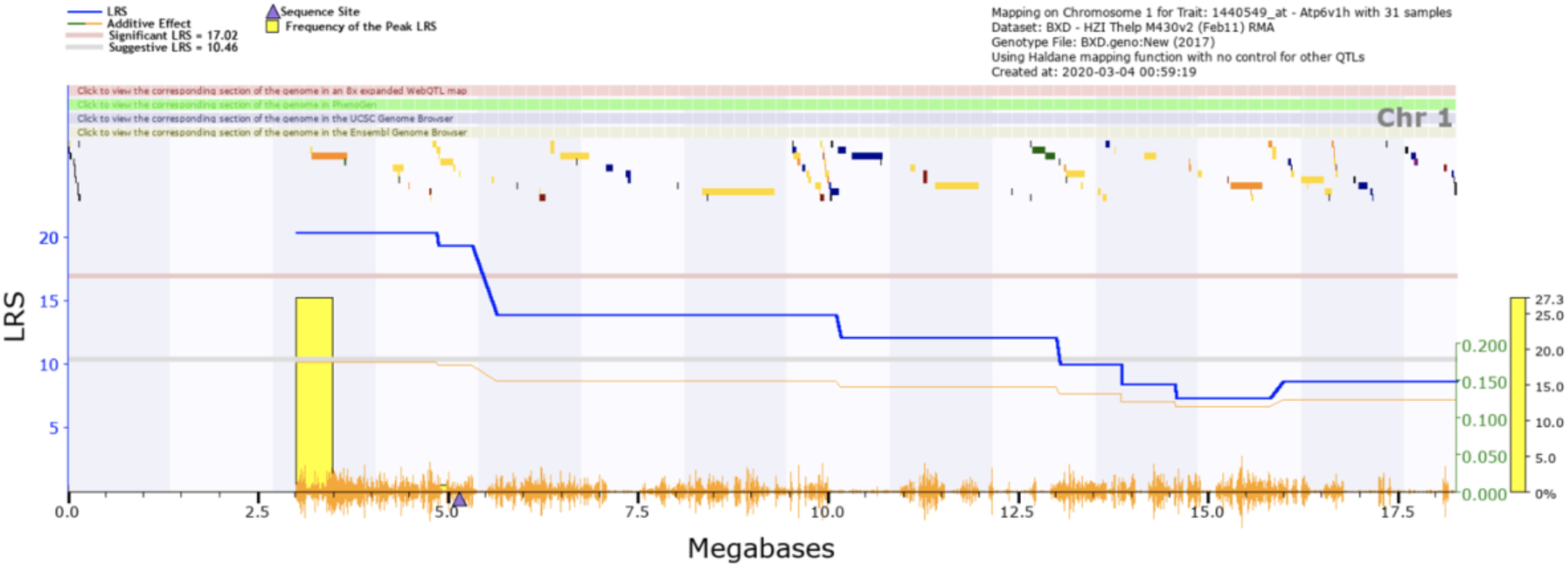
Individual genes and SNPs are shown with the LRS scores shown in blue. The purple triangle is the location of our gene of interest, *Atp6v1h*. The blue line is the LRS value for individual regions on the chromosome. The yellow bar is the frequency of the highest LRS values across the genome. The orange hashes on the bottom by the megabase values are to denote areas of high SNP density.

**Figure 19:**
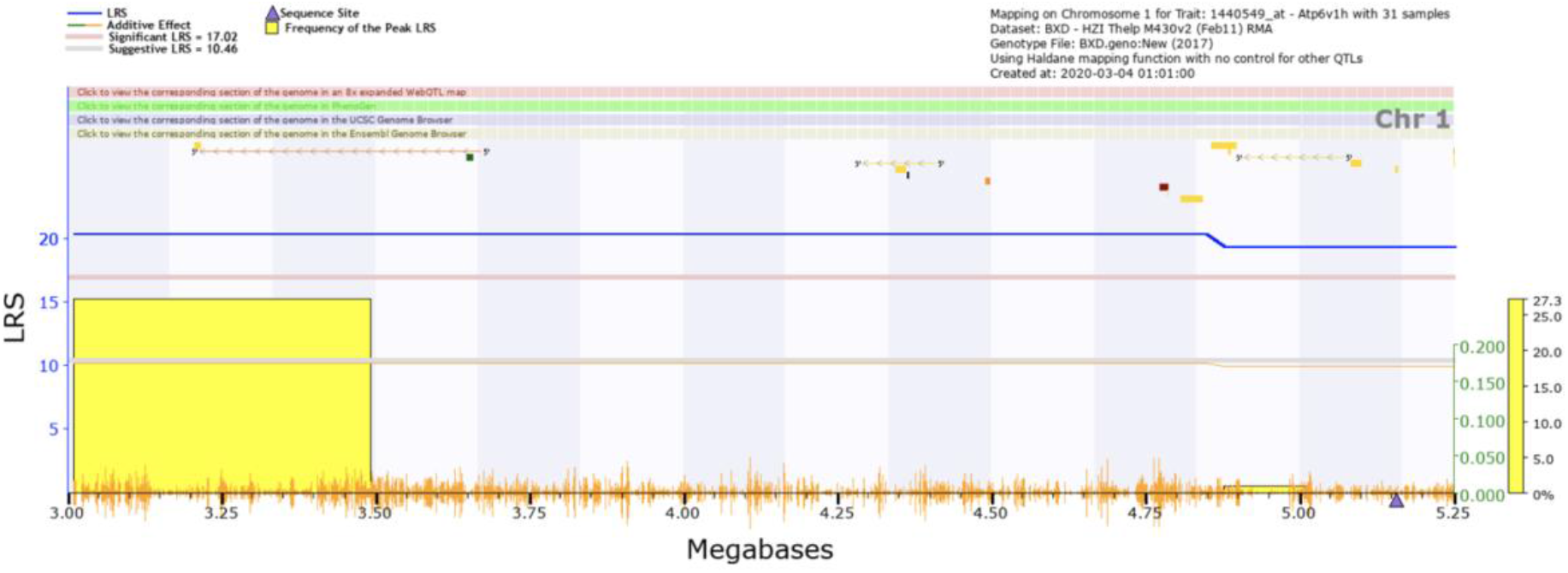
The gene that makes the most sense for this LRS peak is the gene of interest, *Atp6v1h*, which is indicated by the purple triangle at the bottom of the figure.

##### *Trans-*eQTL

In some cases, a gene can be influenced by the expression of another gene at another locus, and whether a gene is *cis*- or *trans*- regulated may be dependent upon the tissue tested, or on environmental factors. To examine this, let us return to *Atp6v1h*, but in a different tissue, and investigate genes that could be influencing *Atp6v1h* expression.

To start the search, change the type of data to liver mRNA (Figure **20A**), the mRNA dataset (Figure **20B**), and enter the specific gene in the search row (Figure **20C**). Once again this will populate a table of records that match the specific parameters of the search. Where we can choose a record of interest, for example based on high LRS score and QTL position, and then verify the probes and normalize the data as demonstrated before (**Figure 21**).

**Figure 20:**
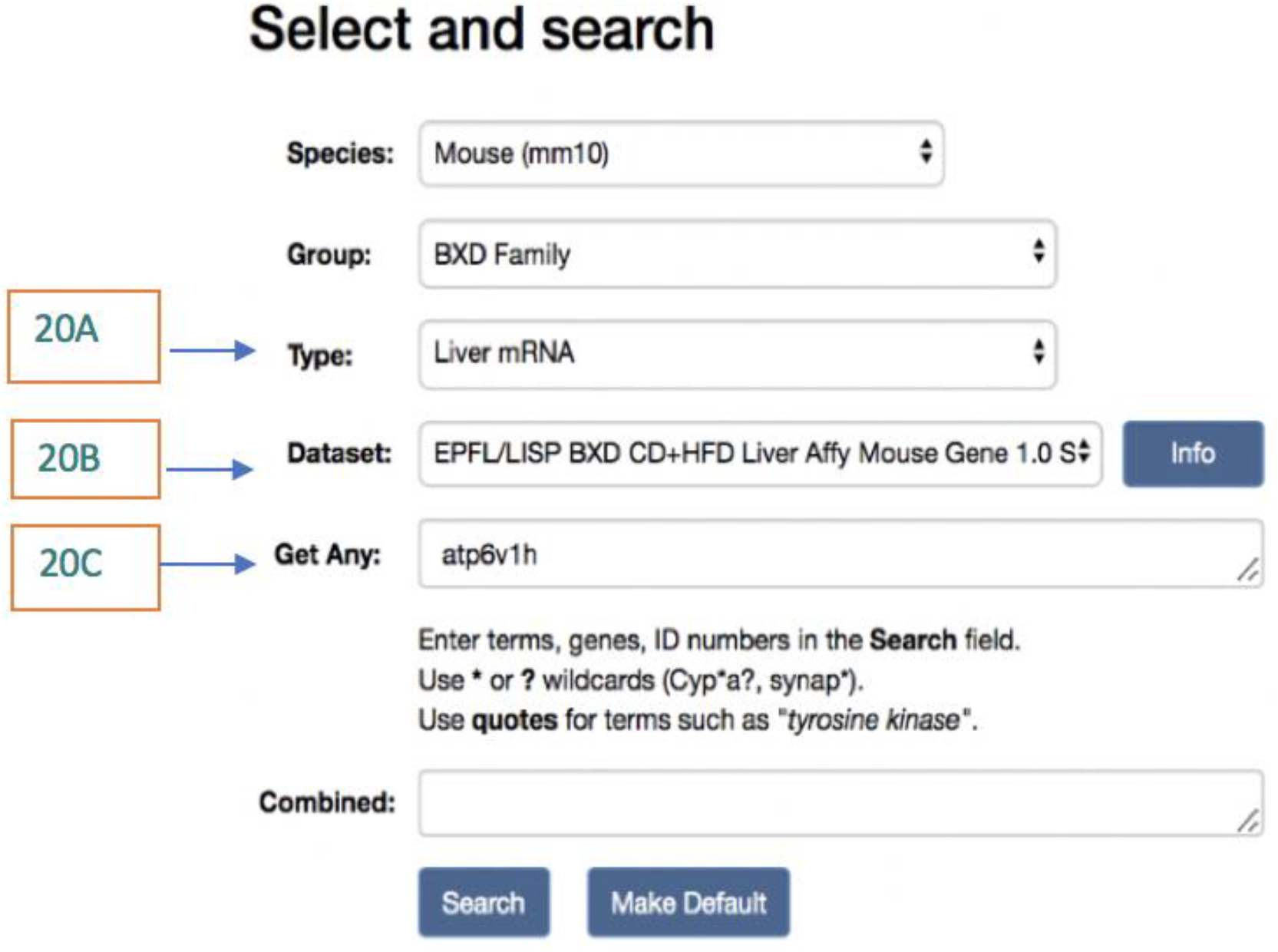
The search parameters for a *trans*-eQTL are similar to the ones for a *cis*-eQTL. For this investigation, we are choosing a different tissue (20A). Make sure to select mRNA datasets that are applicable (20B), and put the gene of interest in the search area (20C).

**Figure 21:**
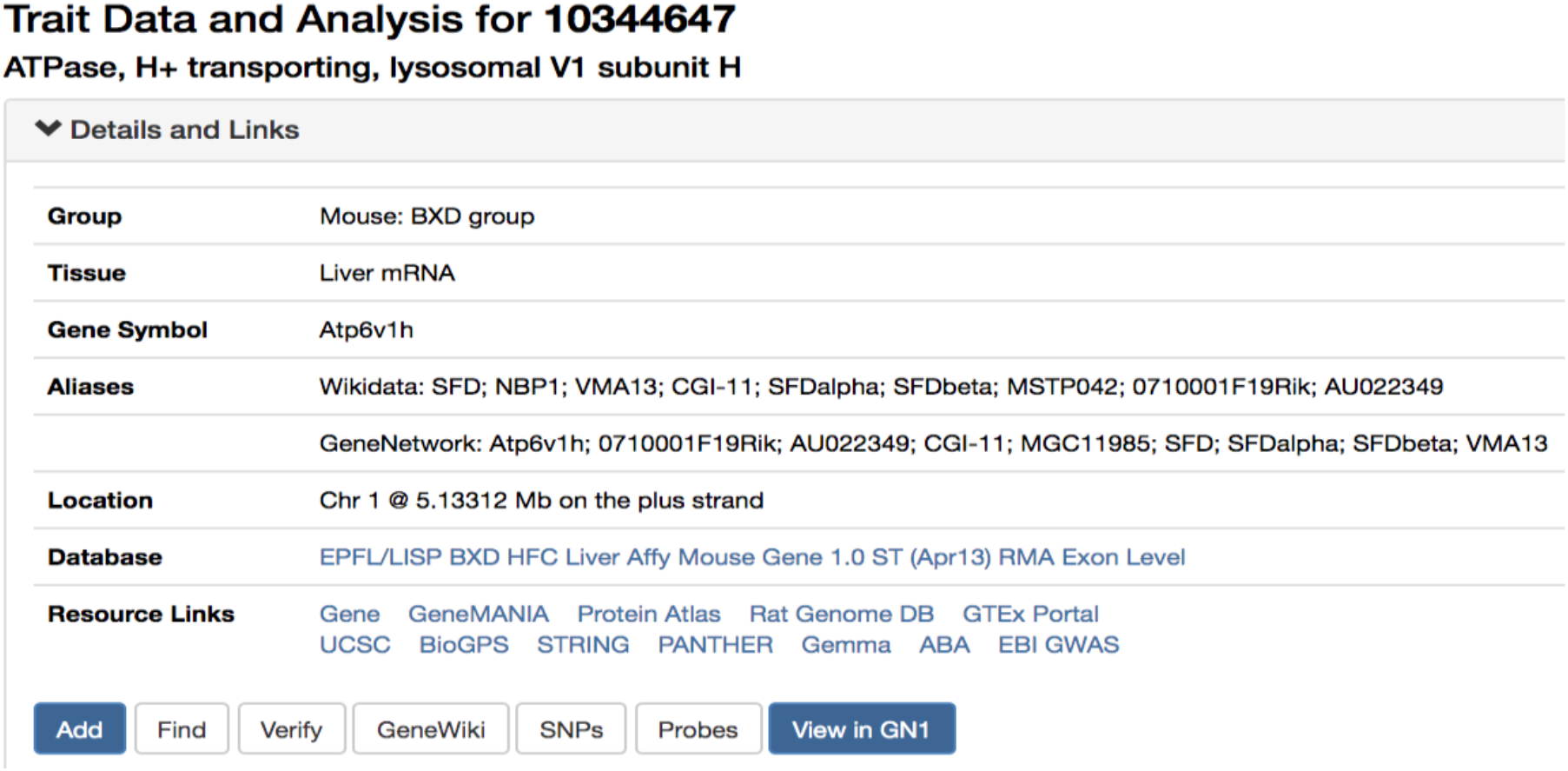
The record with the highest and most significant LRS value in the liver mRNA data set is also a good example of a *trans*-eQTL.

As noted previously, the gene *Atp6v1h* is found on chromosome 1. However, when looking at the mRNA dataset from the liver (Wu et al., 2014), the highest LRS score is found on chromosome 3. This peak surpasses the suggestive LRS value and just reaches significance, as shown in **Figure 22**.

**Figure 22:**
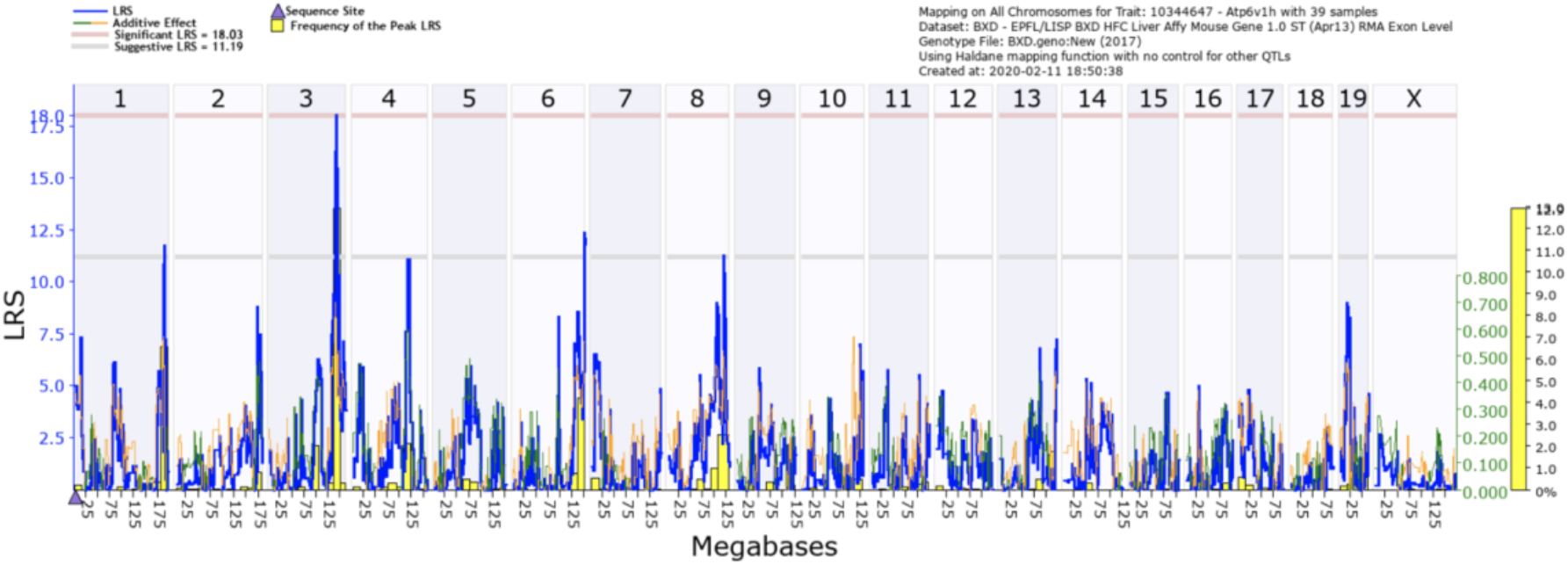
The purple triangle on chromosome 1 shows the location of the gene, *Atp6v1h*. However, the highest, most significant peak is on chromosome 3. This makes this a good candidate for a *trans*-eQTL.

As with the last two case studies, clicking on the chromosome will pull up a map of the genes and SNPs within that chromosome. It is then possible to expand upon that search and narrow the list of genes to only those that are within the QTL confidence interval. **Figure 23** shows the zoomed in view of chromosome 3 with a table of the genes that are located under the peak in **Figure 24**.

**Figure 23:**
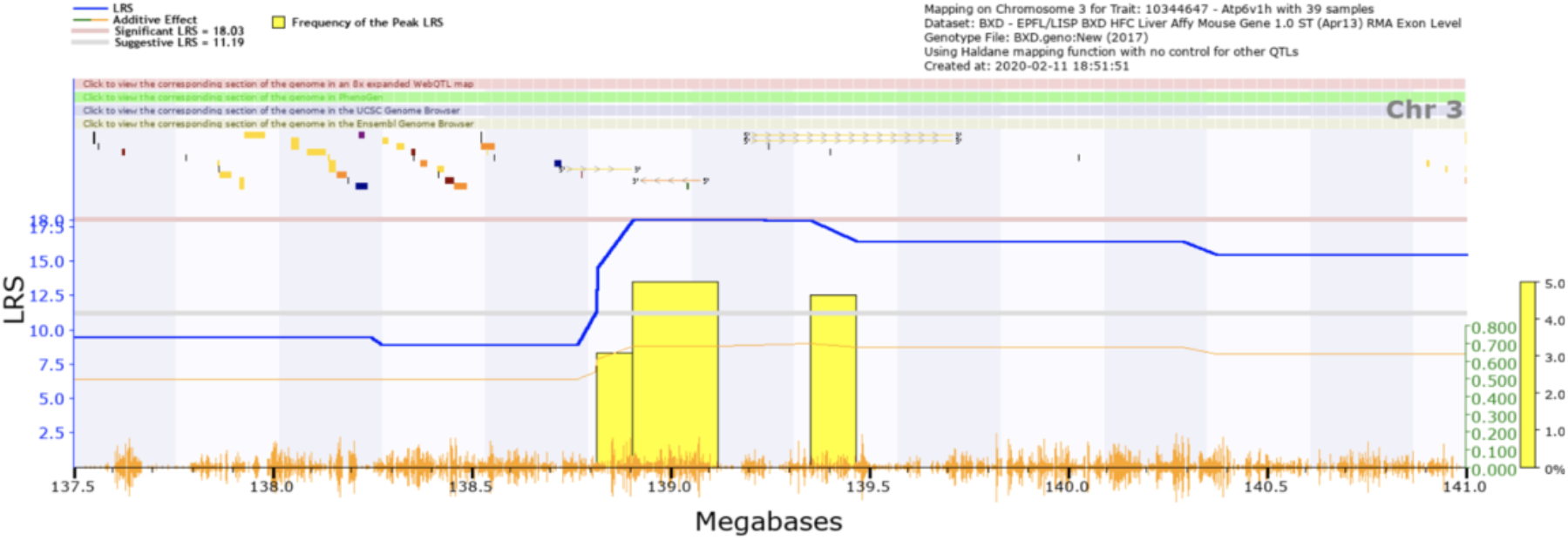
Expanding the map allows for ease of access in the investigation for loci. Since this peak occurs within a *trans*-eQTL region, the purple triangle representing the gene’s location is missing. The blue line is the LRS value for individual regions on the chromosome. The yellow bar is the frequency of the highest LRS values across the genome. The orange hashes on the bottom by the megabase values are to denote areas of high SNP density.

**Figure 24:**
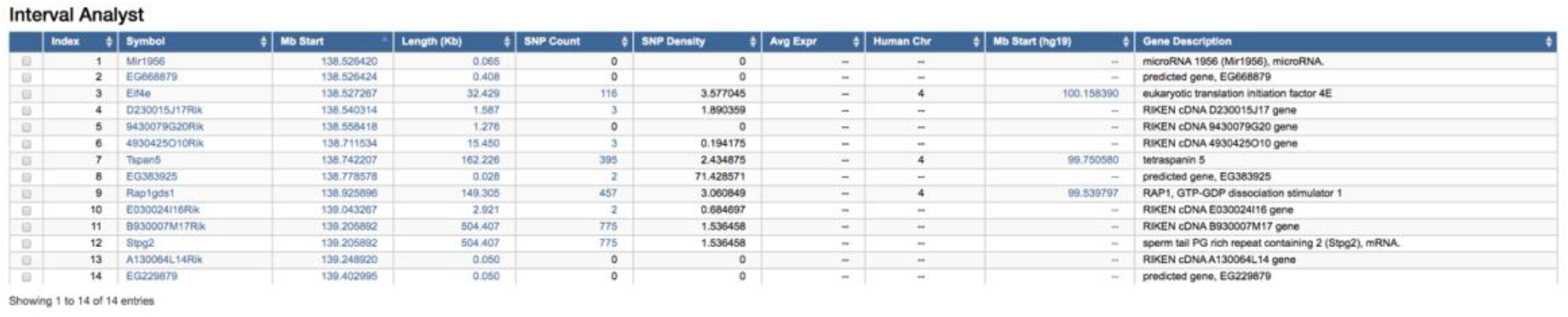
The Interval Analyst table allows researchers to easily investigate genes within the area of interest. This table shows the genes within the area of the LRS peak that could be involved in the trans-regulation of the gene, *Atp6v1h.*

A candidate gene was found based upon the map data and the genes listed within the table. This gene can then be searched in the NCBI database for its genetic information and functions. This is shown in **Figure 25**. This suggests that in liver *Atp6v1h* may be under the control of *Eif4e*, unlike in T helper cells, where it is under its own control. To gain more information, let us look at another immune related tissue, the spleen.

**Figure 25:**
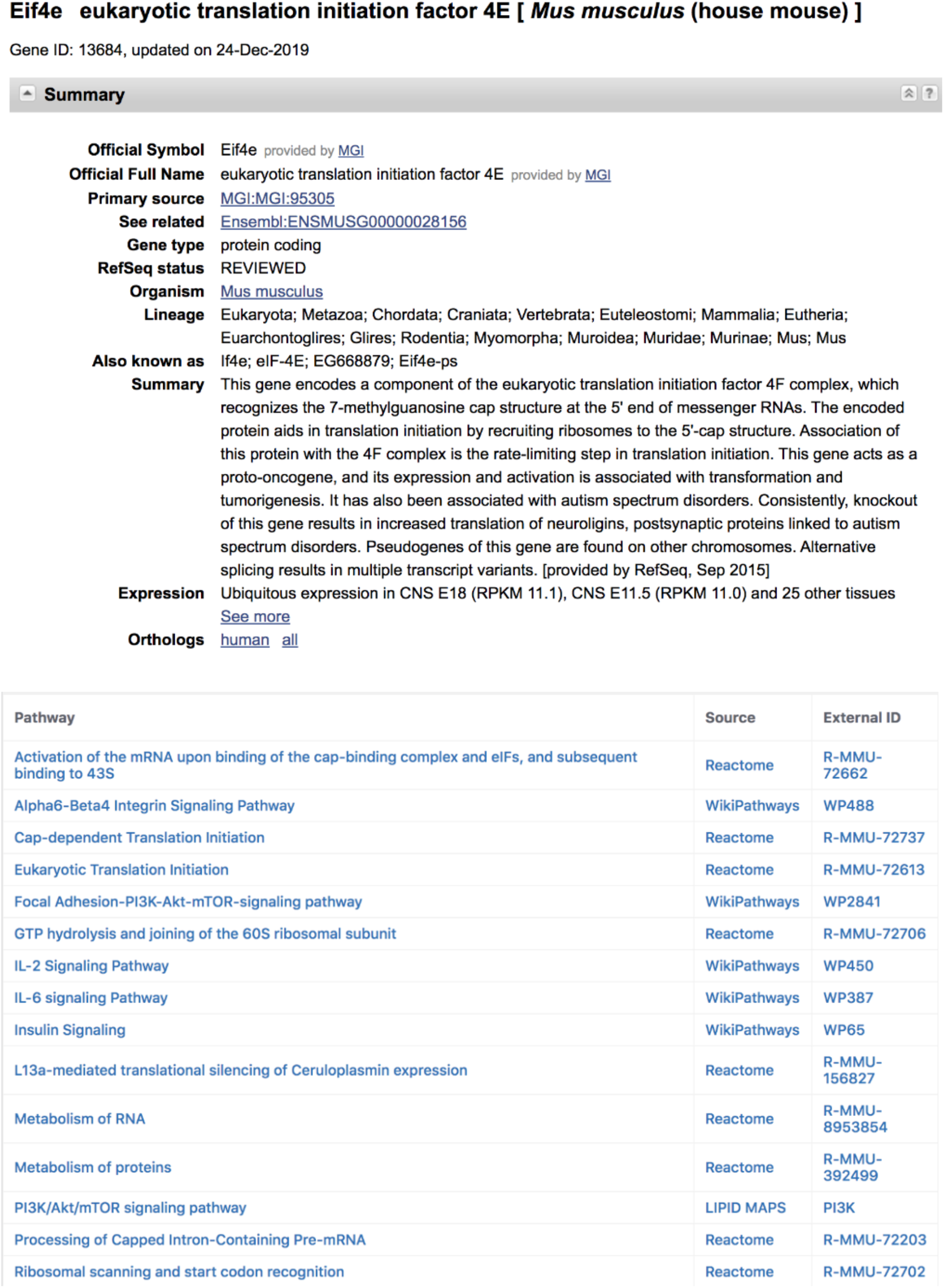
The gene that relates the most with the eQTL shown in Figures 19–22 is *Eif4e*. The genetic summary and functions solidify its role as a regulator of immune responses.

Return to the GN homepage and select ‘Spleen mRNA’ in the ***Type*** box to get gene expression datasets measured in the spleen (**Figure 26A**). Now, when we look under ***Dataset***, we see that there are a number of different datasets which have been collected in spleen, using different technologies and at different institutions. Information about each dataset can be found by clicking the ***Info*** button, for example how the data was analyzed. For the purpose of this example, we will select the exon dataset (UTHSC Affy MoGene 1.0 ST Spleen (Dec10) RMA Exon Level), as an opportunity to examine expression of specific exons of the gene (**Figure 26B**). Again, we search for *Atp6v1h* (**Figure 26C**), and find that this array has 15 probes for *Atp6v1h*, only three of which have significant eQTL, and all three significant eQTL are in *cis*-regulation (**Figure 28**).

**Figure 26:**
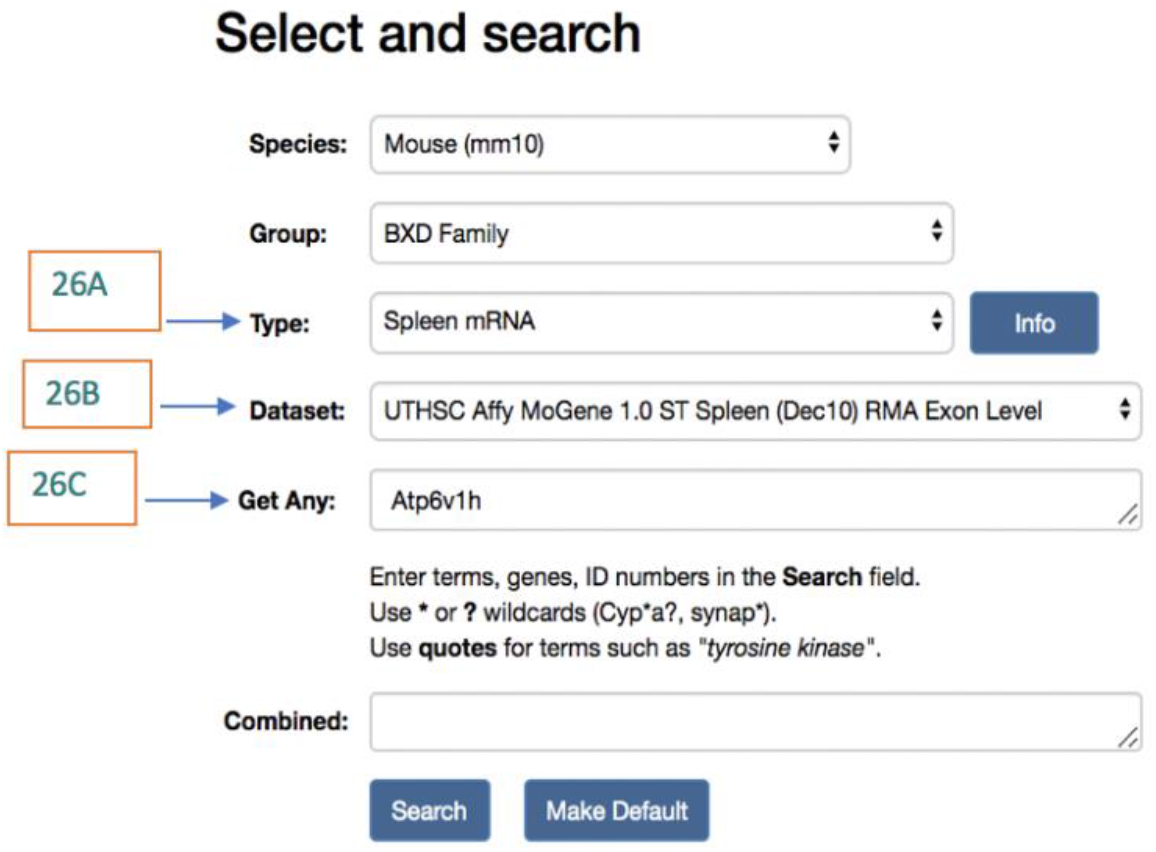
The Select and Search menu in GN. To perform this search, select Spleen mRNA from the *Type* box (**26A**). Within the *Dataset* choose the UTHSC Affymetrix exon level data set (**26B**), and list the gene of interest in the *Get Any* box.

In this case, rather than selecting any one probe, we will select all three (**Figure 27A**), and perform a correlation analysis (**Figures 27B and 28**). This shows us that these three probes are highly correlated, and we can also see the first three principal components. Principal component analysis (PCA) is a technique for data reduction and pattern recognition. The PCA generates eigenvectors that capture the majority of the variation in expression between the traits (Carter, 2013). An eigenvector for gene expression is often referred to as an eigengene, and since these three probes share the same QTL position, this eigengene represents their shared regulation by the Chr 1 QTL.

**Figure 27:**
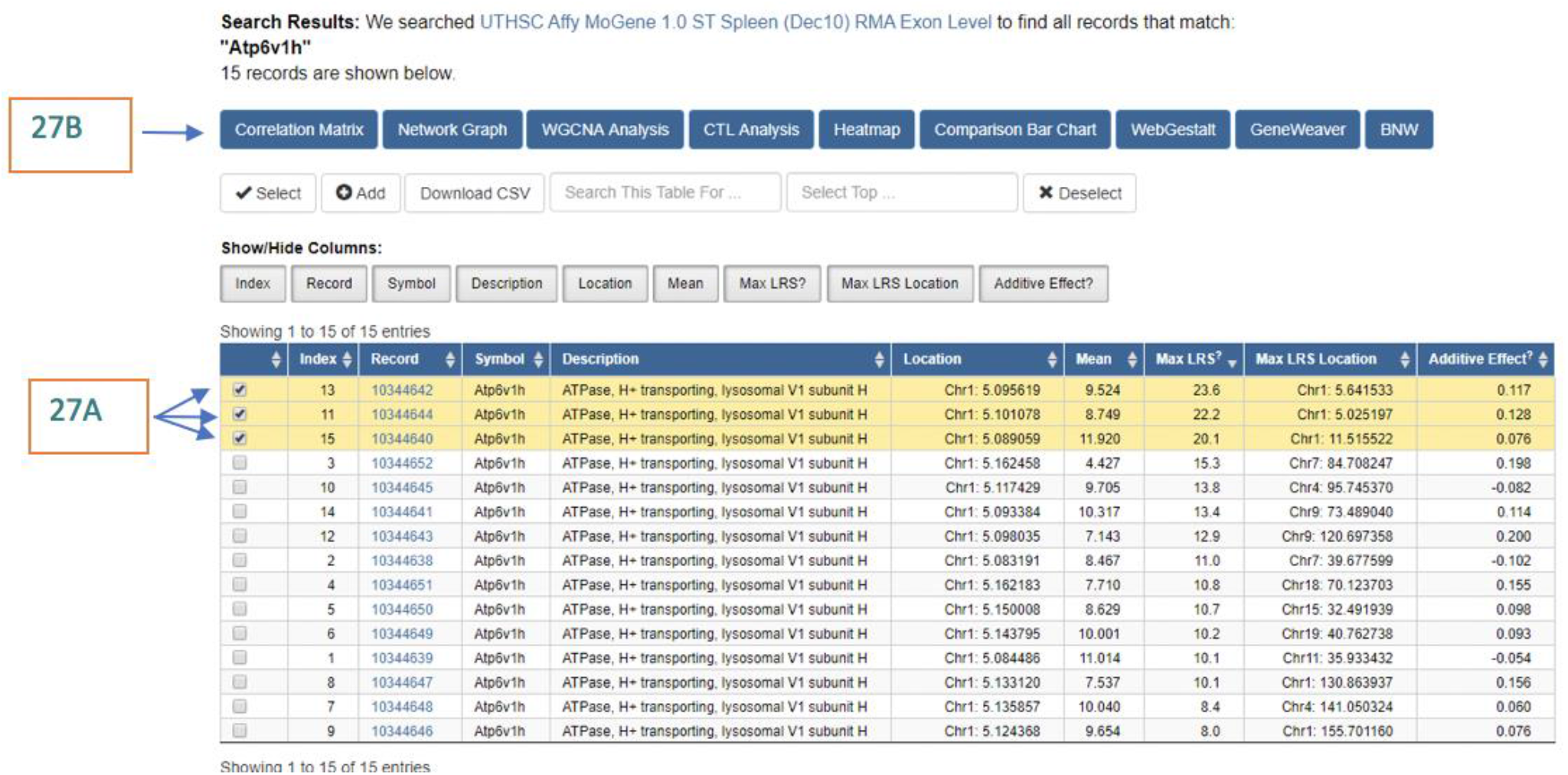
The search results produce a records table that lists the record ID, the gene symbol for the probe, a description of the gene and its location. All of the results are for probes that target the gene of interest, *Atp6v1h*. Of those, only 3 show a significant *cis-*eQTL (10344642, 10344644, and 10344640). For this, we will select all 3 of the probes (**A**), and perform a correlation matrix (**B**).

**Figure28:**
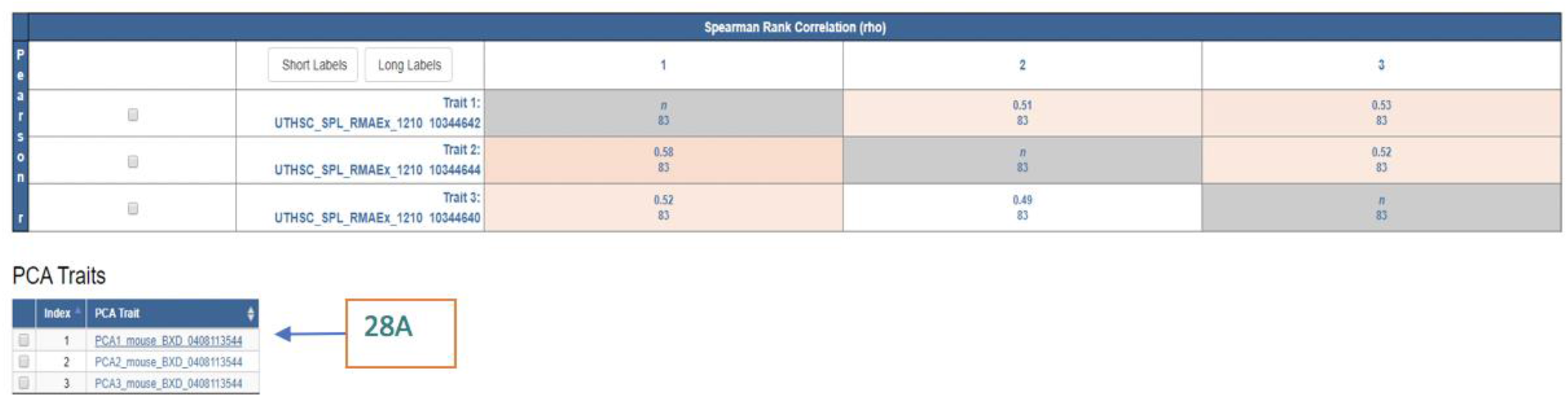
The Correlation Matrix output. Here we can see the PCA traits, also called eigengenes. To further investigate and map an eigengene, select the trait record ID (28A).

This eigengene can be used as a trait in GN by clicking on it (**Figure 28A**). Remapping this trait shows a much stronger peak LRS than any of the probes individually (LRS = 32.53), demonstrating that the eigengene is capturing their shared genetic regulation. This can often help to produce a smaller confidence interval, narrowing the list of potential candidate variants. Given that *Atp6v1h* sits within this interval, this supports *Atp6v1h* having a *cis*-eQTL in spleen.

### 3. Network analyses

We now have two QTL, and we have picked potentially interesting genes within each, but now we want to build up more evidence for which gene in our QTL interval is causal. The first, and most obvious way, is to see what genes our trait of interest correlates with, in tissues that we expect to be related to the trait. We calculated the Spearman’s correlation between the trait BXD_17850 and all probes with expression data in T helper cells (GN319). To do this, go to the Calculate Correlations tab (**Figure 29**), choose the database of interest, and the type of correlation (Pearson, Spearman or Biweight midcorrelation). The user may note that Spearman correlation was used, rather than Pearson’s. The Spearman correlation is considered more powerful when the number of samples is low, which is often the case when comparing between datasets in GN (for example, only 7 strains were used in both BXD_17850 and the T-helper cell expression dataset).

**Figure 29:**
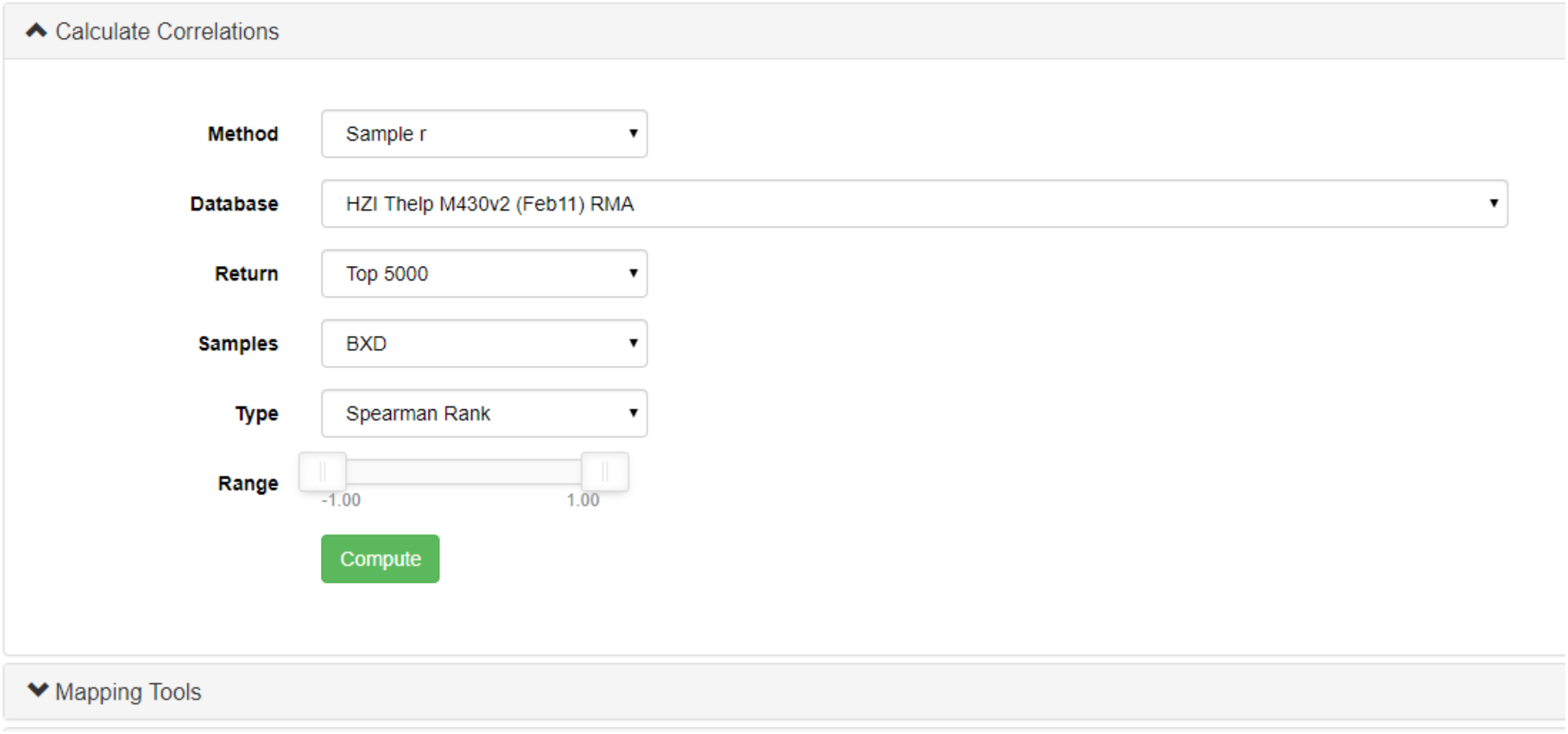
The Calculate Correlations tab on the trait page. The method of correlation, the database for the trait to be correlated against, how many correlations should be returned, the samples to be used, the type of correlation to be used, and the range of correlations to be returned.

Given the high correlations seen, we will take only the most highly correlated genes (rho > 0.85 or < −0.85). To do this, click on the ***More Options*** button (**Figure 30A**), and then change the correlation values, and click on the ***Select Trait*** button (**Figure 30B**). This will reduce the list of correlated probes to only those that meet the new criteria. We now have a list of genes, the expression of which is strongly correlated with our trait. This suggests that these genes are involved with our trait. Next, we want to analyze any enrichment within this gene list. To do this, click ***Select All*** (**Figure 30C**), and then ***WebGestalt*** (**Figure 30D**). WebGestalt is a web client that allows enrichment analysis, and Entrez gene IDs will be directly imported into WebGestalt from GN (Zhang et al., 2005; Wang et al., 2013, 2017; Liao et al., 2019). We used WebGestalt to look for enrichment in Gene Ontology Biological Processes (Ashburner et al., 2000; Harris et al., 2004; Gene Ontology Consortium, 2015; The Gene Ontology Consortium, 2019), KEGG pathways (Kanehisa and Goto, 2000; Kanehisa et al., 2012), Panther pathways (Mi et al., 2019a, 2019b), Reactome pathways (Sidiropoulos et al., 2017; Jassal et al., 2020), and Wikipathway pathways (Pico et al., 2008; Slenter et al., 2018) (**Figure 31**). As many different annotations as wanted can be chosen by clicking on the ‘+’ icon (**Figure 31**). Also note, that the user can change the reference set to match the microarray used to gather the data, if known.

**Figure 30:**
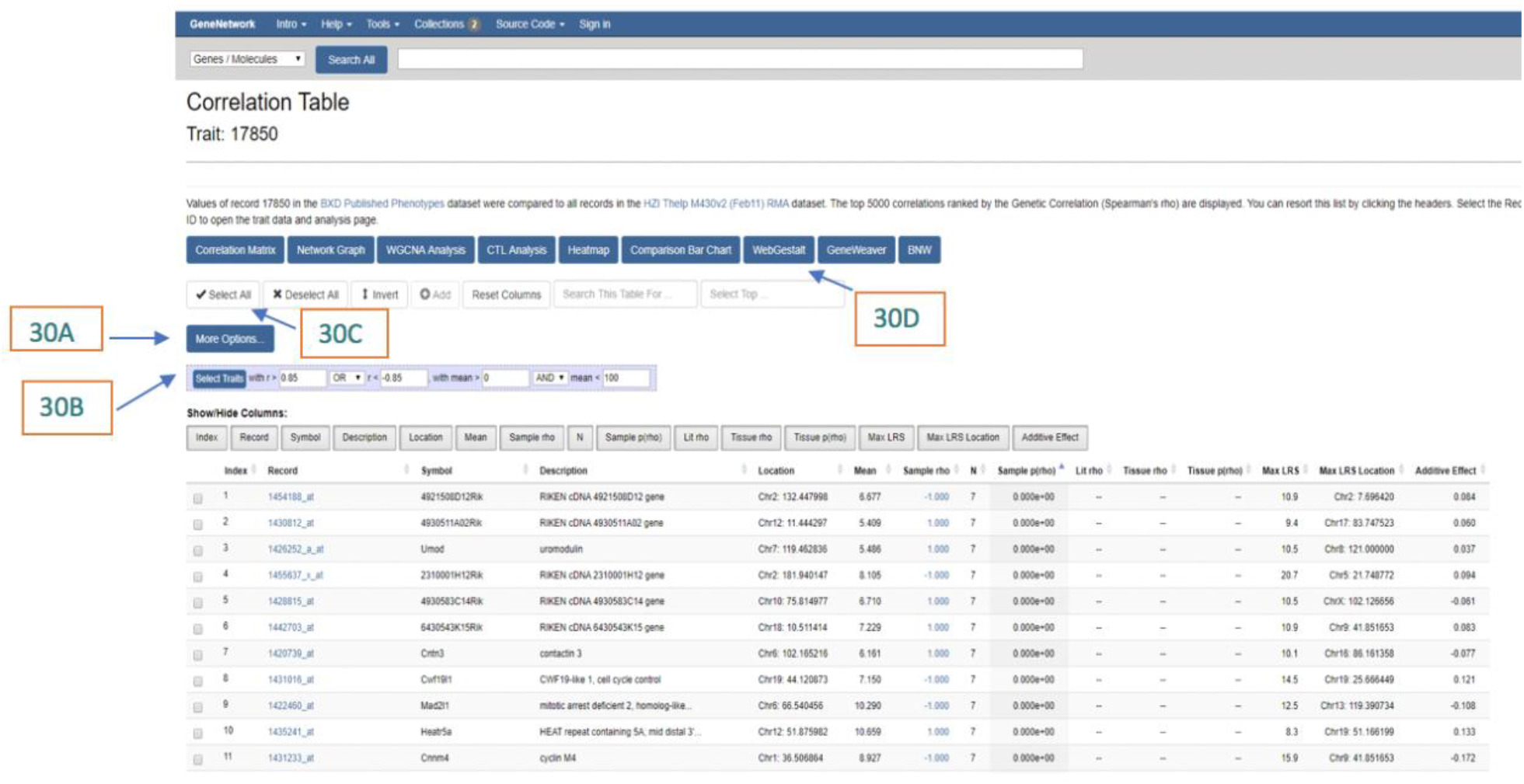
Correlation Table for the eigengene selected. Some useful tools for this search are the *More Options* (**30A**) which will help screen the records table for the most highly correlated genes, the *Select Trait* button (**30B**) allows users to select specific traits, the *Select All* button (**30C**) allows users to select all of the traits in the table, and the *WebGestalt* button (**30D**) opens the window for the WebGestalt search engine.

**Figure 31:**
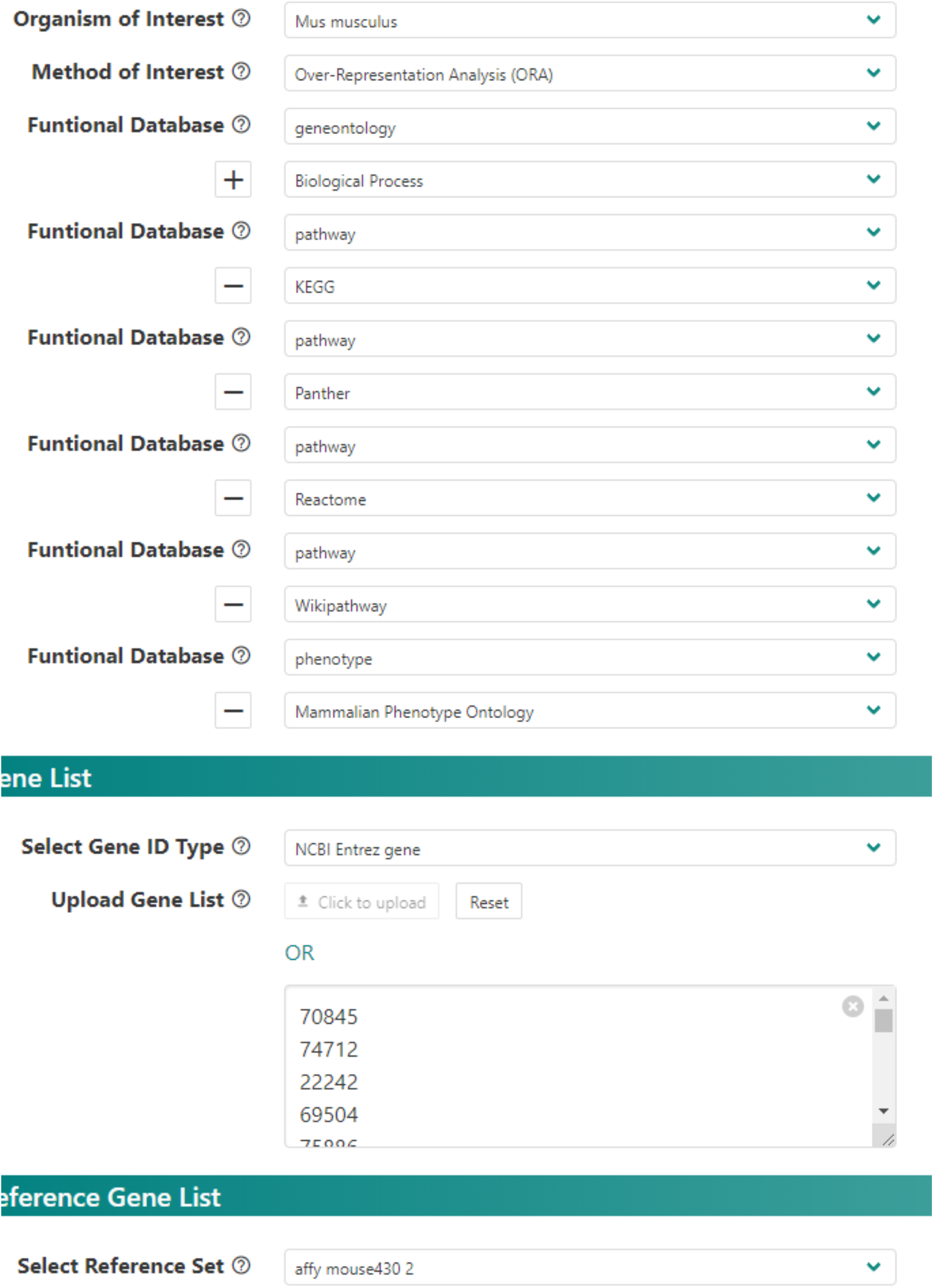
Different functional pathways on WebGestalt. We used WebGestalt to look for enrichment in Gene Ontology Biological Processes, KEGG pathways, Panther pathways, Reactome pathways and Wikipathway pathways.

Click on the submit button at the bottom of the screen to get the enriched annotations. Interestingly, the pathway with the highest enrichment ratio (22.101) was the *lectin pathway of complement activation* (R-MMU-166662), which is a component of the innate immune system (**Figure 32**) (Ali et al., 2012). As we have seen by assessing different tissues, components of this lectin pathway of complement activation are present in all immune related tissues of the body, but are made mostly in the liver (Holers et al., 2020). The lectin pathway of complement activation is present during normal physiology in low levels to monitor for pathogens, present during sickness to recruit macrophages to the sick tissues, and present in autoimmune functions (Farrar et al., 2016).

**Figure 32:**
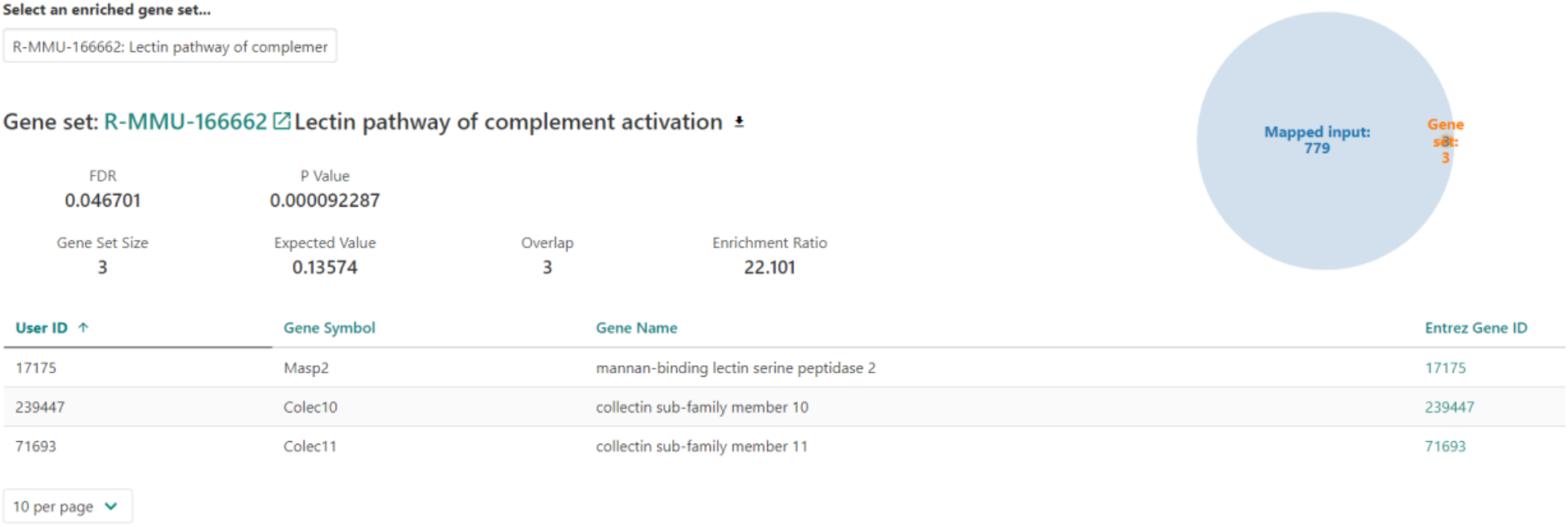
WebGestalt enriched gene set summary. The pathway with the highest enrichment ration was the lectin pathway of complement activation. This pathway is one of three complement pathways that serve to activate macrophages upon invasion from a host.

Returning to the correlation screen (**Figure 30**), we can next narrow the number of genes by looking which of the correlated genes also have suggestive eQTL mapping to our QTL for BXD_17850. To do this, click on the arrow next to the ***Max LRS Location*** header, and then look for the probes with eQTLs within the trait 1.5 LOD interval (i.e. chr1:0-16.107 Mb, observed earlier).

Three genes have strong *cis-*eQTL, our previous candidate *Atp6v1h, Hjurp* and *Mrpl15*. Carrying out enrichment analysis on these 39 probes with a suggestive peak eQTL (LRS > 10) overlapping our QTL showed one significantly enriched annotation, *negative regulation of androgen receptor signaling pathway* (GO:0060766; FDR < 0.036; 3 genes, *Igf1*, *Foxp1*, *Hdac1*; Enrichment ratio 129.13). Interestingly, CCL5 suppresses androgen receptor signaling, suggesting that the genes in this pathway may be downstream of our phenotype (i.e. a variant on Chr 1 alters CCL5 secretion, which in turn suppresses androgen receptor signaling).

Next, we carried out the same analysis in a large spleen dataset (GN283). Correlations were lower here, with the most significant correlation being rho −0.721. Due to this lower correlation, we took a more lenient threshold of (rho > 0.5 or < −0.5). This showed a large number of significantly enriched annotations (**Figure 33**). Although the most significantly enriched annotation does not seem relevant (melanosome assembly), the next four do: interleukin-5 secretion (GO:0072603), regulation of interleukin-5 secretion (GO:2000662), ERKs are inactivated (R-MMU-202670) and Signal Transduction of S1P Receptor (WP57). All of these annotations are involved in the innate immune system, specifically to regulate chemokine (CCL5) and cytokine (IL5) production to increase inflammation, promote macrophage activity, and clear pathogens in all the tissues (Trenchevska et al., 2015; Pydi et al., 2019). Again, we looked for genes which have a peak LRS mapped to the QTL region, and there were no significantly enriched annotations (FDR < 0.05). However, there were eight genes within the QTL region which have *cis*-eQTL, *Mrpl15*, *Scn2a1*, *St18*, *Adhfe1*, *Sntg1, Acvrl1, Snord87 and Eya1*. Interestingly, we see *Mrpl15* again, but not our earlier candidate *Atp6v1h*.

**Figure 33:**
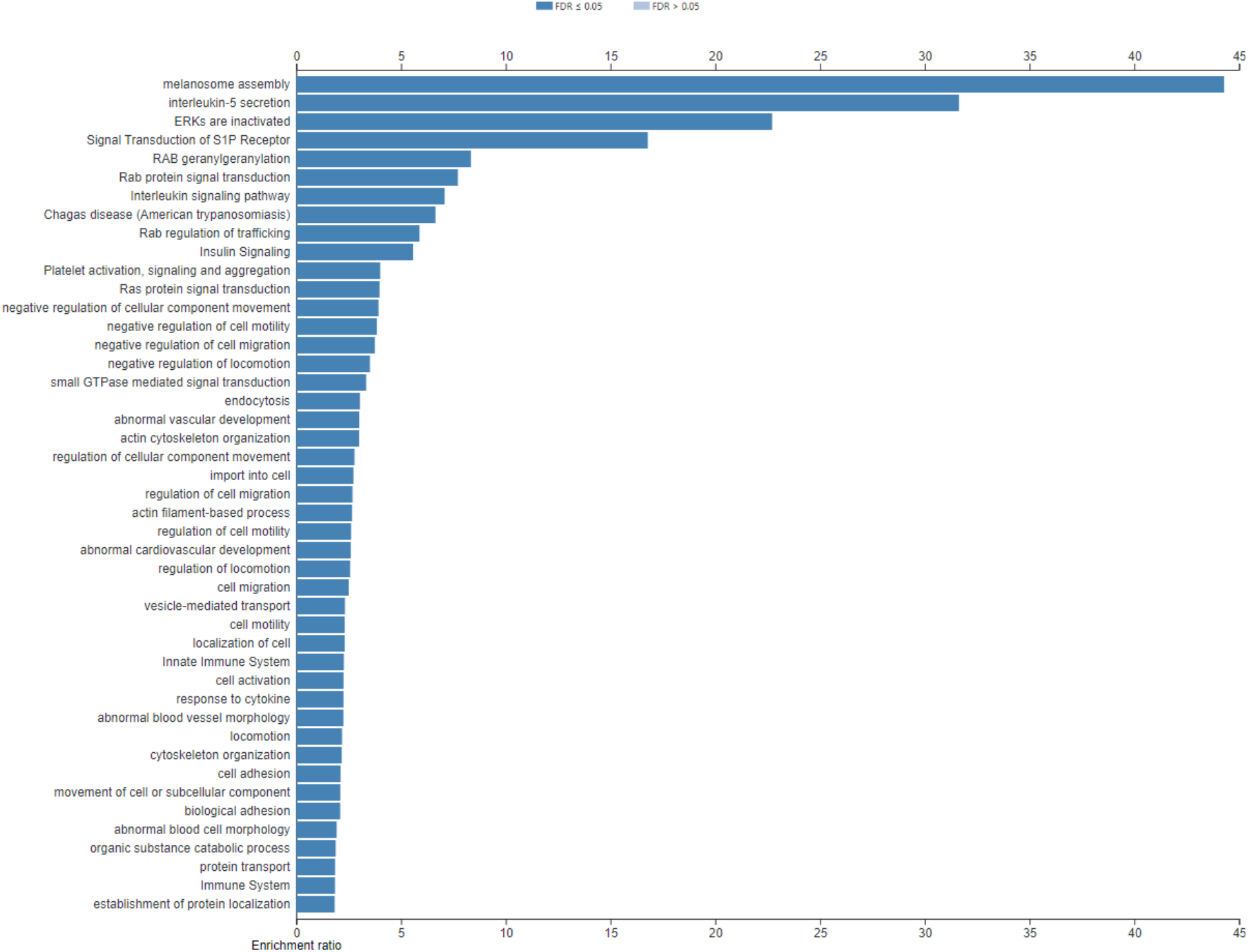
The most significantly enriched annotations associated with the correlation from **Figure 30.** All of these annotations are involved in the innate immune system, specifically to regulate chemokine (CCL5) and cytokine (IL5) production to increase inflammation, promote macrophage activity, and clear pathogens in all the tissues.

So, where do we stand? We have good evidence that CCL5 in the plasma after 12 hours of fasting (trait BXD_17850) is influenced by a QTL on chr1 0-16.107 Mb, and have identified correlated genes which are involved in pathways upstream and downstream of this. We have 8 potential candidates which share an eQTL location, and the expression of which correlate with the phenotype.

We can next correlate the expression of these 8 genes with the whole phenome database in GN, to identify if they may be associated with other immune related phenotypes. To do this, we used the 8 probes and did a correlation with the whole phenome, in a similar way to **Figure 29**, but instead choosing ***BXD published phenotypes***. These were then narrowed down to those with a rho > |0.5|, and a nominal p-value < 0.05. The total number of correlated phenotypes meeting these criteria, and the number of phenotypes which were immune related, were counted, so for each probe we could calculate a percentage of correlated phenotypes which are immune related (**Table 1**).

**Table 1:**
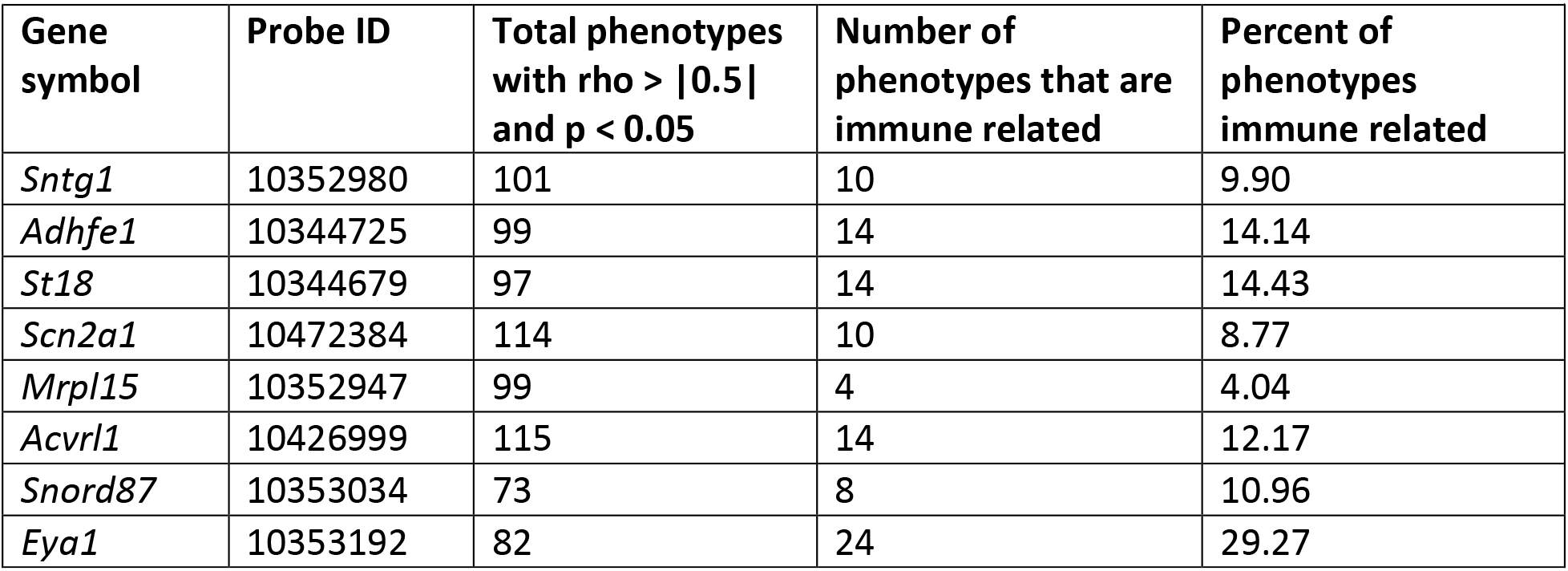
Correlations between spleen gene expression for genes with cis-eQTL in the trait QTL interval, and phenotypes related to immune system. Assumption that causative gene will correlate with a number of immune phenotypes

This suggests that *Eya1* expression is correlated with the highest number of other immune related phenotypes, making it a good candidate. However, four other genes (*Adhfe1*, *St18*, *Acvrl1* and *Snord87*) also correlate with immune phenotypes. Most notably of those, *Acvrl1* and *St18* which are both involved in cytokine release in the immune system. *Acvrl1* signaling is responsible for dendritic cell development within the cytotoxic T cell (CD8α^+^) subtype (Verma et al., 2016). This signaling is also related to inflammatory pathways, similarly to RANTES. *St18* is overexpressed in inflammatory autoimmune diseases and leads to promotion of autoantibodies and inflammation, but also a decrease in the ERK signaling pathway seen in Figure 33 (Radeva et al., 2019).

At this stage, additional analyses could be done in other populations (e.g. if a QTL for the same trait has been detected in another population), or another species (e.g. if there is a GWAS for the phenotype in humans; (Houtkooper et al., 2013; Williams and Auwerx, 2015). These cross-population analyses can often identify a single, top candidate (Ashbrook et al., 2014a, 2015b; Jha et al., 2018b, 2018a; Koutnikova et al., 2009). Alternatively, with the handful of candidates identified, it is practical to move to ‘wet lab’ assays, for example seeing if over- or under-expression of our candidate genes *in vitro* leads to changes in CCL5 levels.

## Conclusion

GeneNetwork is an excellent tool for exploring complex phenotypes with systems genetics. Here we have used GeneNetwork to explore an inflammatory phenotype, and identified a small number of plausible candidate genes. A similar workflow can be used for any trait on GeneNetwork, or for any phenotype collected by an investigator in a genetically diverse population. GeneNetwork can allow users to study relationships between genes, pathways, and phenotypes in an easy to use format.

## Acknowledgments

The authors would like to acknowledge the operators, coders, and contributors to GeneNetwork.org including Robert W. Williams, Pjotr Prins, Saunak Sen, Zachary Sloan, Arthur Centeno, Christian Fischer, Bonface Munyoki, Danny Arends, Sam Ockman, Lei Yan, Xiaodong Zhou, Christian Fernandez, Ning Liu, Rudi Alberts, Elissa Chesler, Sujoy Roy, Evan G. Williams, Alexander G. Williams, Kenneth Manly, and Jintao Wang.

The GeneNetwork site is supported by the University of Tennessee Center for Integrative and Translational Genomics, NIGMS Systems Genetics and Precision Medicine Project (R01 GM123489, 2017-2021), NIDA Core Center of Excellence in Transcriptomics, Systems Genetics, and the Addictome (P30 DA044223, 2017-2022), NIA Translational Systems Genetics of Mitochondria, Metabolism, and Aging (R01AG043930, 2013-2018), NIAAA Integrative Neuroscience Initiative on Alcoholism (U01 AA016662, U01 AA013499, U24 AA013513, U01 AA014425, 2006-2017) NIDA, NIMH, and NIAAA (P20-DA 21131, 2001-2012), NCI MMHCC (U01CA105417), NCRR, BIRN, (U24 RR021760).

## Notes

### Competing Interest Statement

The authors have declared no competing interest.

